# NEMP1 organizes the meiotic telomere–LINC interface to control chromosome movement and telomere integrity

**DOI:** 10.64898/2026.09.17.752236

**Authors:** Hui Zhang, Ling Zhang, Stephanie A. Pangas, Andrea Jurisicova, Helen McNeill

**Author notes:** Corresponding author: Helen McNeill.

## Abstract

Nuclear Envelope Membrane Protein 1 (NEMP1) is highly expressed in oocytes and required for fertility, yet its meiotic function remains poorly understood. Here, we identify NEMP1 as a critical organizer of meiotic telomere–nuclear envelope coupling and chromosome dynamics. Loss of NEMP1 causes telomere aggregation, loss of shelterin protection, telomere shortening, persistent DNA damage, aberrant non-homologous end joining, and chromosome end-to-end fusion, ultimately leading to aneuploidy in mouse oocyte. NEMP1 is required during early fetal meiotic prophase I for telomere–nuclear envelope attachment, bouquet formation, homolog pairing, and synapsis. Live-cell imaging revealed robust rapid prophase movements in wild-type meiocytes, which were nearly abolished by NEMP1 loss and restored by NEMP1-GFP re expression. Mechanistically, NEMP1 associates with telomeric DNA and the SUN1–KASH5 LINC machinery, and SUN1-GFP restores chromosome movement in Nemp1-deficient meiocytes. Together, these findings establish NEMP1 as a nuclear-envelope organizer linking telomere protection, chromosome dynamics, and genome integrity during mammalian oogenesis.

## Introduction

Genome integrity in germ cells is essential for fertility and species continuity^1^. During meiosis, this requirement is challenged by extensive chromosome movement, programmed DNA double strand breaks (DSBs), and prolonged cell cycle arrest, particularly in female germ cells^2–7^. Telomeres play a central role in coordinating these events by anchoring chromosome ends to the inner nuclear membrane, facilitating homolog pairing, synapsis, and recombination during meiotic prophase I^8–12^. Failure to maintain telomere integrity or telomere–NE attachment results in chromosome mis-segregation, aneuploidy, and infertility ^13–15^.

In mammals, meiotic telomere attachment is mediated by a complex and partially redundant network of proteins. Core components include the shelterin complex at telomeres, the meiosis specific TERB1–TERB2–MAJIN complex, and the linker of nucleokinetic and cytoskeleton (LINC) complex containing SUN-domain proteins SUN1 and SUN2 ^9–11,16^. Genetic disruption of TERB or SUN1 impairs telomere–NE tethering and chromosome movement, leading to defective synapsis and sterility^9,17–19^. However, no mammalian model has been described in which meiotic telomere attachment is completely abolished, suggesting the existence of additional, yet unidentified, regulatory factors at the inner nuclear membrane (INM).

Beyond their role in chromosome dynamics, telomeres must be protected from being recognized as DNA breaks ^20^. Critically short or uncapped telomeres activate ATMor ATR dependent DNA damage responses and can be processed by non-homologous end joining (NHEJ), resulting in chromosome end-to-end fusions^21–23^. The shelterin complex suppresses these responses, in part by antagonizing the recruitment of 53BP1, a key mediator of classical NHEJ ^24–27^. In somatic cells, telomere dysfunction leads to genome instability, while in germ cells it has profound consequences for meiotic progression and oocyte quality^28,29^.

Recent work has revealed that the nuclear envelope is not merely a passive boundary but an active platform for genome organization and DNA repair ^30,31^. DNA breaks and dysfunctional telomeres can relocalize to the nuclear periphery, where repair pathway choice and chromatin mobility are regulated through interactions with LINC complexes, nuclear lamina, and cytoskeletal forces ^25,32,33^. These findings raise the possibility that INM proteins may directly participate in telomere protection and DNA damage suppression during meiosis.

NEMP1 is a conserved INM protein and highly expressed in oocytes ^34–36^. Female mice lacking *Nemp1* are severely subfertility, and our previous work has implicated NEMP1 in nuclear envelope integrity and maintain the ovarian reserve ^34,37^. However, how NEMP1 contributes to genome stability in oocytes remains unknown.

Here, we identify NEMP1 as a critical regulator of telomere integrity in oocytes. We found that NEMP1 is highly expressed and enriched at the oocyte nuclear envelope, where it directly associates with telomeres during meiotic prophase, supporting a role for NEMP1 in coordinating telomere–nuclear envelope interactions. Loss of NEMP1 leads to telomere aggregation, shelterin loss, telomere shortening, chromosome end fusion, and aneuploidy in fully grown oocytes. Moreover, during early meiosis, NEMP1 is required to maintain telomere attachment to the NE and to protect meiotic telomeres from ATM–CHK2 activation and 53BP1-mediated NHEJ. Mechanistically, NEMP1 forms a functional complex with SUN1 and shelterin, linking telomeric DNA to the INM. Together, our findings reveal NEMP1 as a previously unrecognized nuclear envelope–based telomere guardian essential for female fertility.

## Results

### NEMP1 depletion disrupts meiotic maturation, spindle assembly, and chromosome segregation

To validate NEMP1 depletion in oocytes, we first examined NEMP1 expression by immunofluorescence staining. In wild-type (WT) germinal vesicle (GV) oocytes, NEMP1 was strongly enriched at the nuclear envelope. In contrast, NEMP1 immunofluorescence was completely absent in *Nemp1* knockout GV oocytes, confirming the efficient depletion of NEMP1 (Extended Data Fig. 1a). Consistent with these results, western blot analysis of whole ovaries showed reduced NEMP1 protein levels in heterozygous ovaries and a complete loss of NEMP1 protein in *Nemp1* knockout ovaries (Extended Data Fig. 1b). We previously reported spindle defects in ovulated Nemp1 knockout oocytes ^34^. To further investigate the role of NEMP1 in oocyte meiotic progression, we monitored germinal vesicle breakdown (GVBD) and first polar body extrusion (PBE) during in vitro maturation. NEMP1 depletion did not affect the ovary area and number of oocytes obtained (Extended data Fig. 1 c-f) or the rate of GVBD but markedly reduced the PBE rate (Fig. 1a-d), indicating that NEMP1 is required for normal meiotic maturation. Consistent with the reduced PBE rate, oocytes from knockout females frequently arrested at metaphase I (MI) (Fig 1 e; Extended data Fig. 1 g, Supplementary video 1-2) and exhibited severe spindle assembly defects, including elongated, short, multipolar, and unfocused spindles (Fig. 1f-h; Extended data Fig. 1 h). NEMP1 depletion also resulted in substantial chromosome misalignment at MI (Fig. 1i). Furthermore, analysis of metaphase II (MII) chromosome spreads combined with CentB FISH revealed a significantly increased incidence of aneuploidy in Nemp1 knockout oocytes (Fig. 1j, k). Together, these findings demonstrate that NEMP1 is essential for meiotic spindle assembly and accurate chromosome segregation, thereby maintaining chromosome euploidy during oocyte maturation.

**Figure 1.**
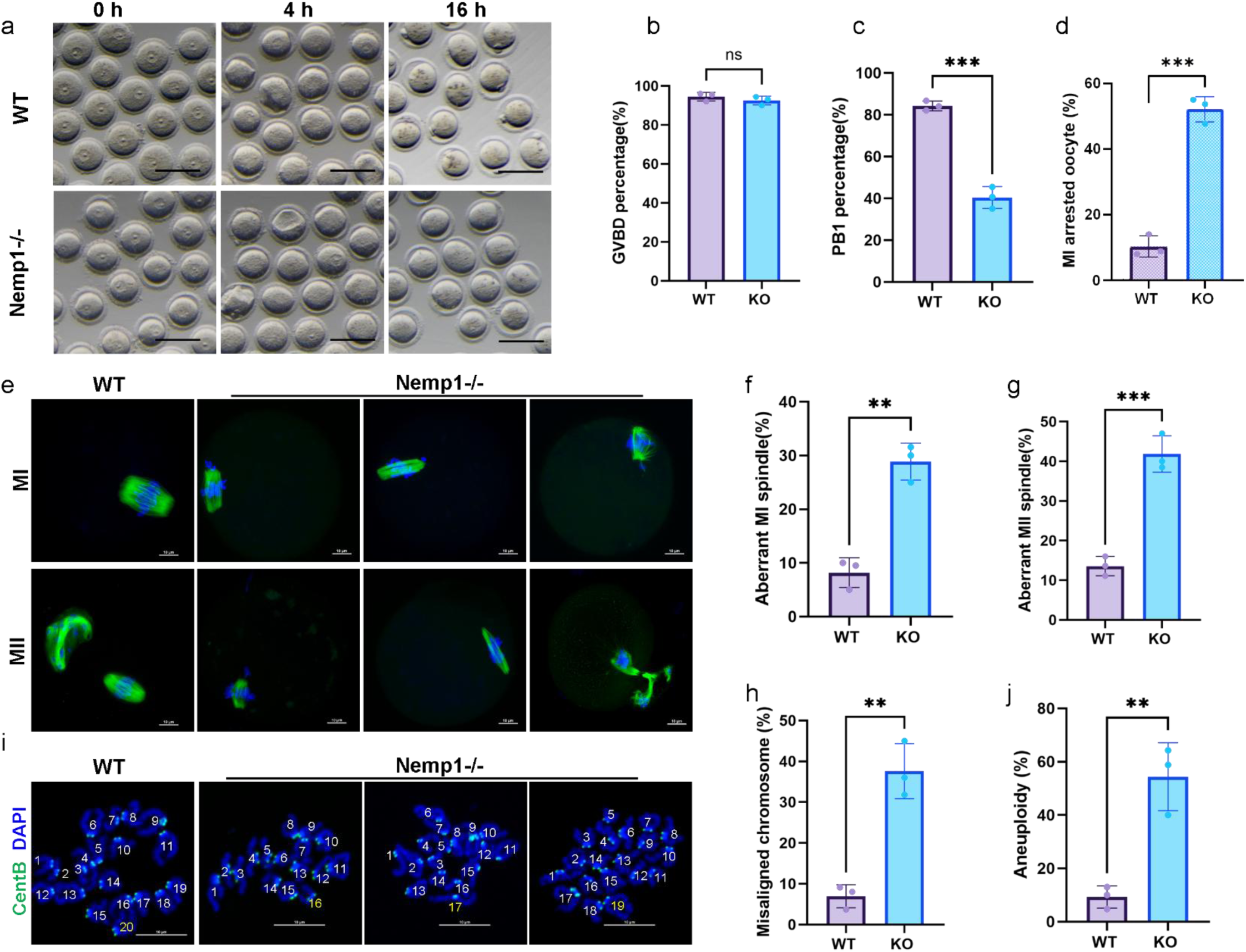
NEMP1 depletion impairs meiotic maturation, spindle assembly, and chromosome segregation in mouse oocytes. (a) Representative images showing germinal vesicle (GV), germinal vesicle breakdown (GVBD), and first polar body extrusion (PBE) during in vitro maturation of control and *Nemp1*-depleted oocytes. Scale bars, 100 μm. (b– d) Quantification of oocyte recovery, GVBD, and PBE rates, respectively. NEMP1 depletion did not affect oocyte recovery or GVBD but significantly reduced the rate of PBE. (e) Representative images of control and *Nemp1*⁻/⁻ oocytes at metaphase I (MI) following in vitro maturation. (f–h) Representative images and quantification of spindle morphology, showing severe spindle assembly defects in *Nemp1*⁻/⁻ oocytes, including elongated, monopolar, multipolar, and unfocused-polar spindles. (i) Representative images and quantification of chromosome alignment defects in MI oocytes following NEMP1 depletion. (j) Representative metaphase II (MII) chromosome spreads and CentB fluorescence in situ hybridization (FISH) images from control and *Nemp1*⁻/⁻ oocytes. (k) Quantification of aneuploidy in MII oocytes, demonstrating a significant increase in chromosome abnormalities following NEMP1 depletion. Data are the mean ± s.d. of n = 3 independent trials per group (≥20 viable oocytes per trial). Statistical analyses were performed using two-sided unpaired t-tests. *P < 0.05, **P < 0.01, ***P < 0.001; ns, not significant. Scale bars, 10 μm in f and J.

### NEMP1 maintains telomere organization and integrity during oocyte development

Aneuploidy is a major form of genomic instability in female germ cells and a leading cause of miscarriage and developmental disorders^38^, while telomere dysfunction may contribute to its development ^39,40^. We next assessed oocyte telomere spatial organization. In control oocytes, telomeres appeared as discrete, well-spaced foci distributed throughout the nucleoplasm. In striking contrast, Nemp1 knockout oocytes exhibited prominent telomere aggregates (TAs), in which multiple telomeres coalesced into large foci (Figure 2 a, Extended Data Fig. 2a; Supplementary Video 3-4), a phenotype previously reported in human tumor cells and senescent mesenchymal stem cells ^41,42^. Quantitative analysis revealed that ∼54.46% of Nemp1 knockout GV oocytes displayed TAs, compared with ∼9.78% in controls (p < 0.0001) (Figure 2 b), demonstrating a highly penetrant telomere disorganization phenotype upon NEMP1 loss. Consistent with this, *Nemp1* knockout oocytes exhibited a significant reduction in the number of telomere foci per nucleus (Figure 2 c). Telomere instability has been associated with intranuclear lamina structures and increased telomere–lamina interactions in cellular senescence models ^42–45^. To determine whether the observed phenotype reflected altered nuclear lamina and NE integrity, we examined lamin A/C levels and distribution in GV oocyte (Extended Data Fig. 2 b, c). No differences were detected between control and *Nemp1* knockout oocytes (Extended Data Fig. 2 c), indicating that telomere instability occurred independently of overt changes in lamina abundance. Importantly, comparable telomere abnormalities were not detected in *Nemp1* knockout mouse embryonic fibroblasts (MEFs) (Extended Data Fig. 2d, e), suggesting that NEMP1 plays a cell type-specific role in maintaining telomere organization and that its requirement for telomere stability may be particularly critical in female germ cells.

**Figure 2.**
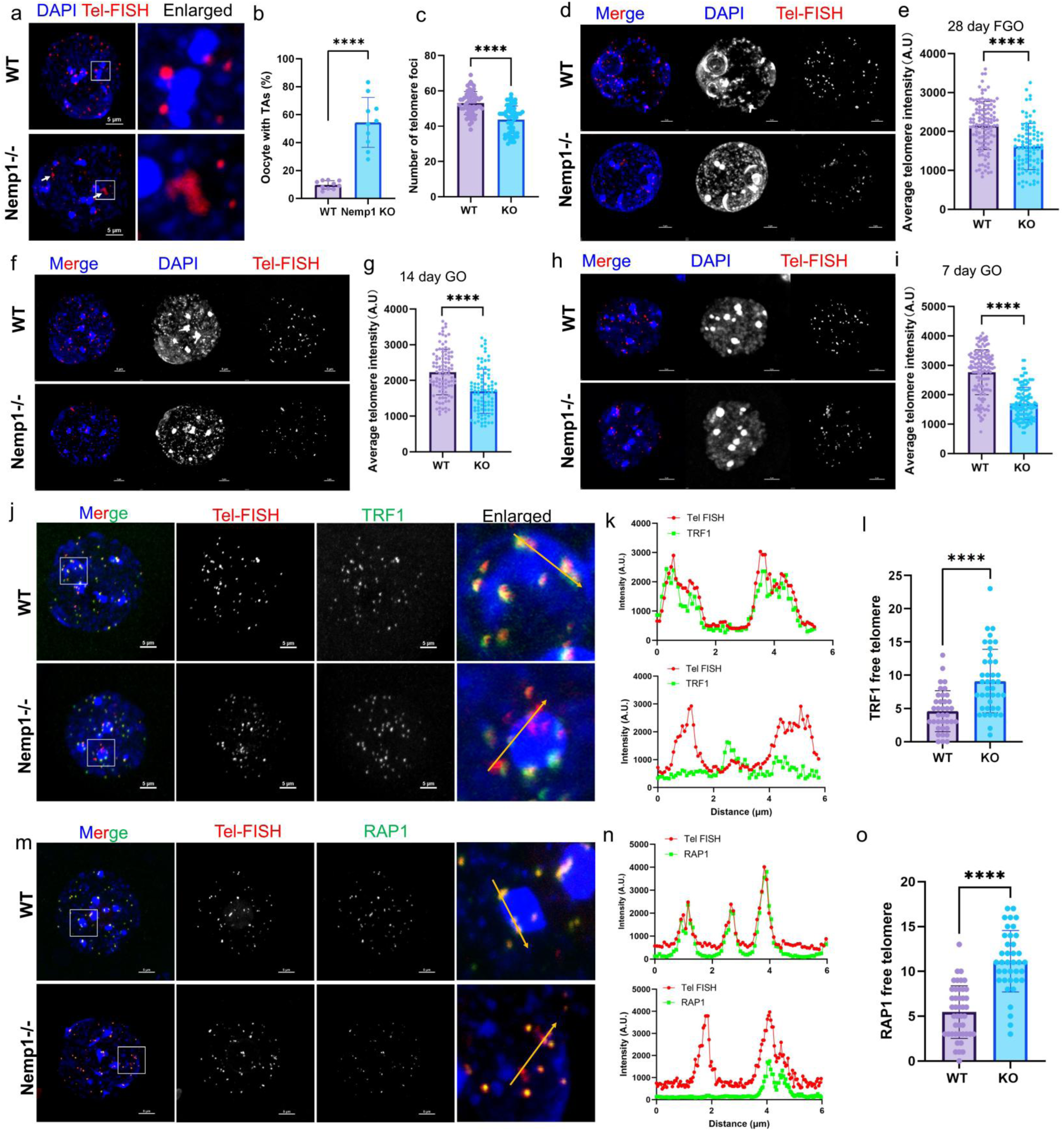
Loss of NEMP1 disrupts telomere organization and promotes telomere aggregation in oocytes. (a) Representative images of telomere FISH in control and *Nemp1*⁻/⁻ germinal vesicle (GV) oocytes. Telomeres are shown as discrete foci in control oocytes but form prominent aggregates (TAs) in *Nemp1*⁻/⁻ oocytes. DNA was counterstained with DAPI. Scale bar, 5μm. (b) Quantification of the percentage of GV oocytes exhibiting telomere aggregates in control and *Nemp1*⁻/⁻ groups. (c) Quantification of the number of telomere foci per nucleus in control and *Nemp1*⁻/⁻ GV oocytes. (d–i) Telomere FISH fluorescence intensity analysis in growing oocytes (GO) and fully grown oocytes (FGO) from control and *Nemp1*⁻/⁻ females. (j) Representative images of GV-stage oocytes stained for telomeres FISH and the shelterin protein TRF1. (k) Fluorescence intensity profiles of TRF1 and telomere FISH along the yellow line in (j) Quantification of TRF1-free telomeres in control and *Nemp1*⁻/⁻ oocytes. (m) Representative images of GV-stage oocytes stained for telomeres FISH and the shelterin protein RAP1. (n) Fluorescence intensity profiles of TRF1 and telomere FISH along the yellow line in (m). (o) Quantification of RAP1-free telomeres in control and *Nemp1*⁻/⁻ oocytes. Data are the mean ± s.d. of n = 3 independent trials per group (≥15 viable oocytes per trial). Statistical analyses were performed using two-sided unpaired t-tests. ****P < 0.0001. Scale bars, 10 μm.

We employed telomere quantitative fluorescence in situ hybridization (Q-FISH) ^46^ in oocyte from control and *Nemp1* knockout females. Average telomere fluorescent intensity was markedly reduced in *Nemp1* knockout oocytes, both in growing oocytes (GO) and fully grown oocytes (FGO), indicating impaired telomere maintenance (Figure 2 d-i). Due to the limited number of oocytes, we assessed telomere length using a newly developed quantitative PCR–based assay that quantifies telomere DNA low-input samples, in which telomeric DNA abundance was normalized to the nuclear reference gene *Rn18s*^47^ (Extended Fig 3 a). Representative amplification curves for telomere DNA and *Rn18s* demonstrated reproducible amplification across samples (Extended Data Fig. 3b–f), with corresponding cycle threshold (Ct) values shown in Extended Data Fig. 3d–g. Using this approach, we found that relative telomere length was significantly reduced in both *Nemp1* knockout GO and FGO compared with wild-type controls (Extended Data Fig. 3h, i). Together, these complementary Q-FISH and qPCR analyses demonstrate that NEMP1 loss is associated with progressive telomere shortening in oocytes.

Given the pronounced TAs and telomere shortening observed in Nemp1 knockout oocytes, we next asked whether NEMP1 is required to maintain telomere protection and shelterin association at chromosome ends. Deletion of Nemp1 in GV oocytes caused a significant increase in “free” telomeres lacking shelterin components, as evidenced by reduced occupancy of the shelterin proteins TRF1 and RAP1 at chromosome ends (Figure 2 j, m, Supplementary Video 5-6). Quantitative analysis of TRF1 and RAP1 foci free telomere revealed markedly diminished shelterin recruitment in *Nemp1* knockout compared with controls (Figure 2 k, l, n and o). Together, these findings demonstrate that loss of NEMP1 leads to altered telomere organization without global disruption of nuclear envelope integrity, supporting a specific role for NEMP1 in maintaining telomere architecture within the oocyte nucleus.

### Loss of NEMP1 disrupts telomere organization and promotes chromosome fusion in maturing oocytes

We next asked whether telomere abnormalities persist during oocyte maturation. Using telomere FISH combined with tubulin immunofluorescence, we found that telomere aggregates (TAs) persisted in both metaphase I (MI) and metaphase II (MII) *Nemp1* knockout oocytes (Figure 3 ad). Because loss of telomere-capping proteins or critically shortened telomeric repeats can cause telomere dysfunction, characterized by telomere aggregation and chromosome end-to-end fusion ^48–50^, we next examined chromosome integrity during meiotic maturation. Analysis of metaphase I (MI) chromosome spreads revealed pronounced chromosome abnormalities in *Nemp1* knockout oocytes, including chromosome bridges, chromosome ends lacking detectable telomeric signals, end-to-end chromosome fusions, and chromosome fragments (Extended Data Fig. 4). Among these abnormalities, end-to-end chromosome fusions (Fig. 3f) and chromosome ends lacking detectable telomeric signals (Fig. 3g) were particularly prominent. These defects were accompanied by a significant reduction in telomere fluorescence intensity (Fig. 3h) and a decrease in the number of detectable telomere foci (Fig. 3i). Such chromosome end-to-end fusions are a hallmark of dysfunctional or deprotected telomeres and are consistent with telomere-driven end joining ^33,49,51,52^.

**Figure 3.**
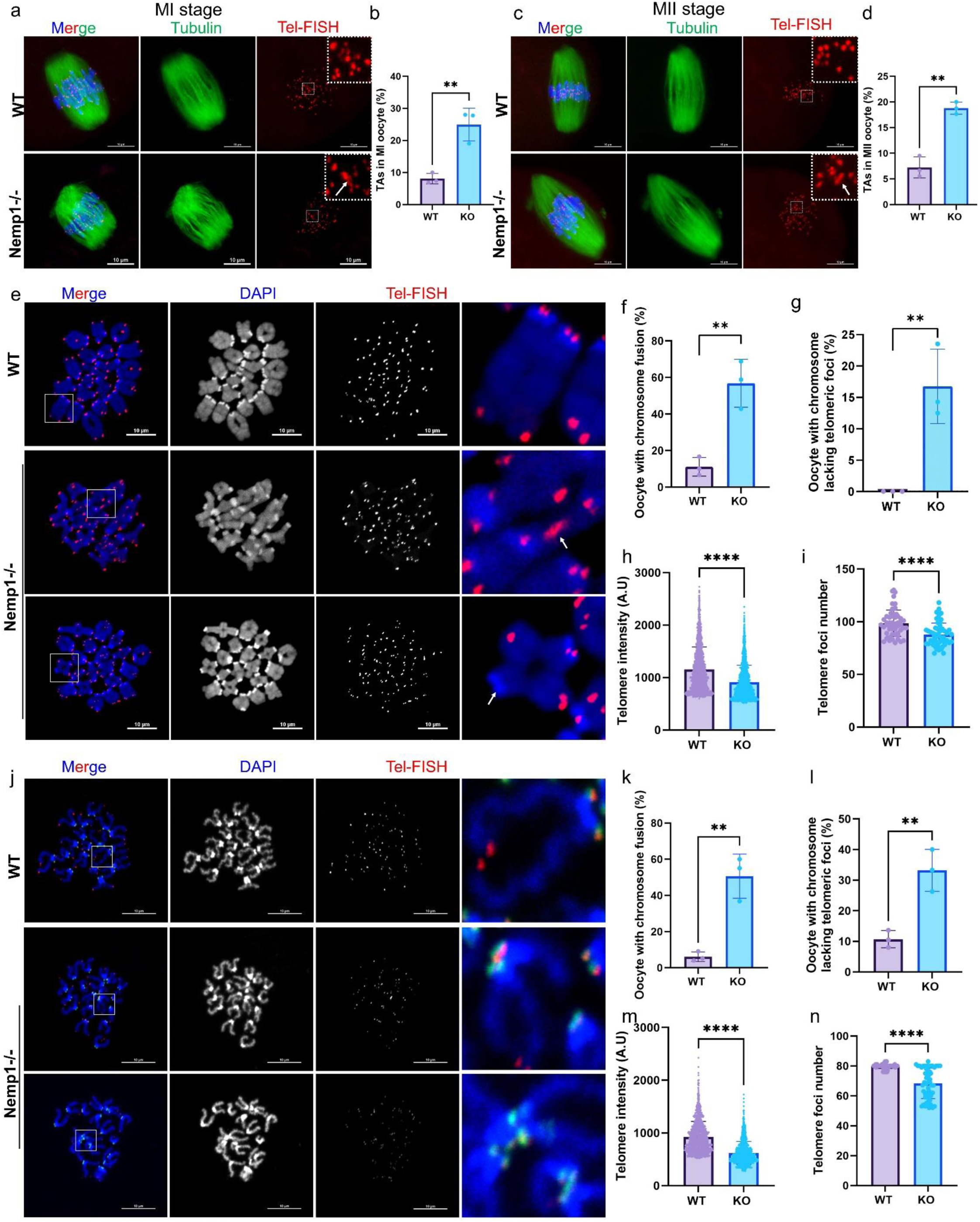
Loss of NEMP1 disrupts telomere organization and promotes chromosome fusion during oocyte maturation. (A) Representative images of metaphase I (MI) and metaphase II (MII) WT and *Nemp1*⁻/⁻ oocytes stained for tubulin and telomere FISH. Telomeres are indicated by telomere FISH signals. Arrowheads indicate telomere aggregates (TAs). (B, D) Quantification of telomere aggregates in MI and MII WT and *Nemp1*⁻/⁻ oocytes, respectively. (C) Representative images of tubulin immunofluorescence and telomere FISH in MII oocytes. (E) Representative MI chromosome spreads from WT and *Nemp1*⁻/⁻ oocytes showing chromosome morphology and telomere FISH signals. Arrows indicate chromosome end-to-end fusions. (F) Quantification of chromosome endto-end fusion frequency in MI oocytes. (G) Quantification of the percentage of chromosomes lacking detectable telomere FISH signals in MI chromosome spreads. (H) Quantification of telomere FISH fluorescence intensity in MI oocytes. (I) Quantification of telomere foci number in MI oocytes. (J) Representative MII chromosome spreads from WT and *Nemp1*⁻/⁻ oocytes showing chromosome morphology and telomere FISH signals. Arrows indicate chromosome end-to-end fusions. (K) Quantification of chromosome end-to-end fusion frequency in MII oocytes. (L) Quantification of the percentage of chromosomes lacking detectable telomere FISH signals in MII chromosome spreads. (M) Quantification of telomere FISH fluorescence intensity in MII oocytes. (N) Quantification of telomere foci number in MII oocytes. Data are the mean ± s.d. of n = 3 independent trials per group (≥15 viable oocytes per trial). Statistical analyses were performed using two-sided unpaired t-tests. **P < 0.01, ****P < 0.0001. Scale bars, 10 μm.

These chromosome abnormalities persisted into meiosis II. Chromosome spreads from MII *Nemp1* knockout oocytes showed a high frequency of chromosome end-to-end fusions (Figure 3j, k), together with an increased proportion of chromosomes lacking detectable telomeric signals (Figure 3l), reduced telomere FISH intensity (Figure 3m), and decreased telomere foci number (Figure 3n). Because chromosome fusions and telomere dysfunction can compromise accurate chromosome segregation and promote aneuploidy, these findings establish a direct link between NEMP1-dependent telomere organization, meiotic chromosome integrity, and the severe subfertility observed in *Nemp1* knockout females. Oocyte aneuploidy is a major cause of embryonic developmental failure and pregnancy loss ^53^, highlighting the potential importance of NEMP1-mediated telomere protection in maintaining oocyte developmental competence.

We next sought to define the developmental window during which NEMP1 establishes and maintains telomere integrity in the female germline. To this end, we conditionally deleted *Nemp1* in growing oocytes using the *Nemp1*^flox/flox^; Zp3-Cre model, in which Cre-mediated recombination occurs after the primary follicle stage (Extended Data Fig. 5a–c). Immunofluorescence confirmed efficient depletion of NEMP1 in oocytes while NEMP1 expression was retained in the surrounding cumulus cells (Extended Data Fig. 5 b, c). Strikingly, unlike constitutive *Nemp1* knockout oocytes, Zp3-Cre–mediated deletion of *Nemp1* did not induce detectable telomere aggregates or alter telomere organization (Extended Data Fig. 5 d, e). Consistent with this finding, telomere fluorescence intensity and relative telomere length were comparable between control and conditional knockout GV oocytes (Extended Data Fig. 5 f, g).

We next asked whether restoring NEMP1 at this later developmental stage could reverse the telomere defects established in the constitutive Nemp1 knockout background. NEMP1-GFP was re-expressed from a Rosa-*Nemp1*-GFP allele under the control of Zp3-Cre (Extended Data Fig. 5 h, i). Despite robust NEMP1-GFP expression, telomere aggregates persisted in *Nemp1* knockout oocytes (Extended Data Fig. 5 j, k), and telomere fluorescence intensity remained reduced (Extended Data Fig. 5 l). Thus, restoring NEMP1 after the primary follicle stage was insufficient to reverse the telomere abnormalities caused by earlier NEMP1 loss.

Together, these findings demonstrate that NEMP1 is essential for maintaining shelterin stability and telomere integrity at chromosome ends, and its telomere-protective function is established during early oogenesis, likely beginning in fetal development.

### NEMP1 is required for telomere anchoring and protection during fetal meiosis

We next investigated whether NEMP1 functions during the early stages of meiosis. During early meiotic prophase I, attachment of telomeres to the nuclear envelope (NE) is essential for homologous synapsis and recombination ^54–56^. *Nemp1* knockout meiocytes exhibited a significant increase in internal telomeres compared with wild-type meiocytes (Figure 4a, b). To directly examine the ultrastructural organization of NE-associated telomeres, we performed electron microscopy (EM) of meiotic telomeres. In wild-type meiocytes, EM revealed characteristic electron-dense conical thickenings at the ends of synapsed lateral elements (LEs), together with a distinct capping structure associated with the inner nuclear membrane (INM) (Figure 4 c), which is consistent with previously reported^9,57^. Strikingly, these structures were frequently disrupted in *Nemp1* knockout meiocytes, including at the ends of synapsed LEs positioned adjacent to the INM (Figure 4 c), indicating a profound disruption of the ultrastructural organization of NEassociated telomeres.

**Figure 4.**
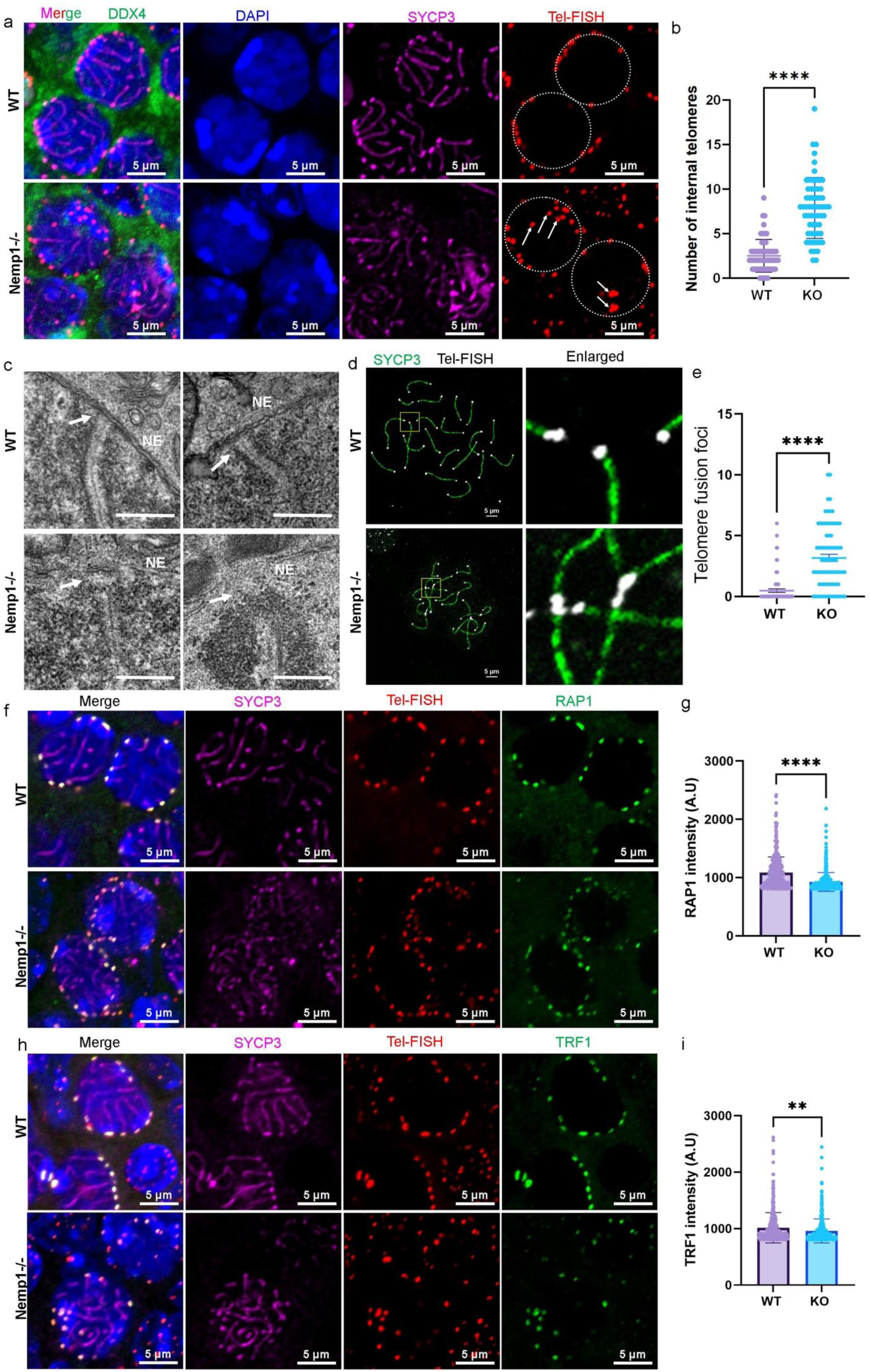
NEMP1 is required for meiotic telomere anchoring and protection during early prophase I. (A) Representative images of pachytene-stage WT and *Nemp1*⁻/⁻ meiocytes showing telomere localization relative to the nuclear envelope (NE). Telomeres were detected by telomere FISH, and the NE was visualized by immunostaining. Arrowheads indicate telomeres detached from the NE. (B) Quantification of the percentage of internalized telomeres in WT and *Nemp1*⁻/⁻ meiocytes. (C) Representative transmission electron microscopy (TEM) images of meiotic telomeres in WT and *Nemp1*⁻/⁻ meiocytes. In wild-type meiocytes, electron-dense conical thickenings at the ends of synapsed lateral elements (LEs) and a telomere-associated capping structure at the inner nuclear membrane (INM) are evident (arrowheads). These structures are absent in WT meiocytes, including at synapsed LE ends adjacent to the INM. (D) Representative pachytene meiocyte spreads stained for SYCP3 and telomere FISH in WT and *Nemp1*⁻/⁻ oocytes. Arrowheads indicate telomere aggregates, and arrows indicate chromosome end-to-end fusions. (E) Quantification of chromosome end-to-end fusions in pachytene-stage meiocytes. (F, H) Representative images of TRF1 and RAP1 localization at telomeres in WT and *Nemp1*⁻/⁻ meiocytes. Telomeres were visualized by FISH. (G, I) Quantification of telomere-associated TRF1 and RAP1 fluorescence intensity in WT and *Nemp1*⁻/⁻ meiocytes. Data are the mean ± s.d. of n = 3 independent ovaries from 3 mice. Statistical analyses were performed using two-sided unpaired t-tests. **P < 0.01, ****P < 0.0001. Scale bars, 5 μm.

We next examined telomere organization and chromosome integrity in pachytene meiocytes using SYCP3 immunostaining combined with telomere FISH (Figure 4 d). *Nemp1* knockout pachytene meiocytes exhibited increased telomere aggregation and a significantly higher frequency of chromosome end-to-end fusions compared with wild-type controls (Figure 4 d, e). These findings indicate that defective telomere anchoring is associated with compromised chromosome-end integrity during early meiotic prophase.

Because shelterin proteins are essential for protecting chromosome ends, we next examined the recruitment of TRF1 and RAP1 to telomeres. Loss of NEMP1 disrupted NE-associated telomere positioning and significantly reduced the accumulation of both TRF1 and RAP1 at telomeres, as evidenced by decreased fluorescence intensity (Figure 4 f–i). Thus, NEMP1 loss compromises not only the physical anchoring of telomeres to the NE but also their association with key shelterin components.

Together, these findings demonstrate that NEMP1 functions during early meiotic prophase in fetal oocytes to establish and maintain NE-associated telomere architecture. Loss of NEMP1 disrupts telomere anchoring and shelterin recruitment, leading to persistent telomere instability and chromosome end-to-end fusion that cannot be corrected during subsequent oocyte maturation.

### NEMP1 prevents accumulation of unrepaired telomeric DNA breaks and inappropriate recombination during meiosis

Telomere deprotection is known to trigger persistent DNA damage responses (DDR) at chromosome ends ^58,58–60^. Our recent research showed that depletion of NEMP1 in fetal and early postnatal ovaries results in robust activation of the DNA damage response ^37^. We therefore asked whether the telomere defects observed in *Nemp1* knockout oocytes are associated with DNA damage signaling during meiotic prophase I. We first examined telomere integrity and DDR activation across meiotic prophase I in E17.5 meiocytes using chromosome spreads stained for telomeric DNA, SYCP3 and γH2AX. Telomere fluorescence intensity was reduced in Nemp1 knockout meiocytes beginning at the zygotene stage and remained reduced through subsequent stages of meiotic prophase I, consistent with impaired telomere maintenance (Extended Data Fig. 6a-f). In parallel, γH2AX signal remained elevated in *Nemp1* knockout meiocytes during the pachytene and diplotene stages, indicating persistent DNA damage signaling during late meiotic prophase I (Extended Data Fig. 6g, h).

We next examined whether this DDR was specifically associated with chromosome ends. Consistent with the telomere shortening and disorganization observed in *Nemp1* knockout oocytes, we detected increased γH2AX and phosphorylated CHK2 at telomeric chromosome ends, particularly in pachytene-stage meiocytes (Fig. 5a–d). These findings indicate that loss of NEMP1 results in telomere-associated DNA damage during meiotic prophase I.

**Figure 5.**
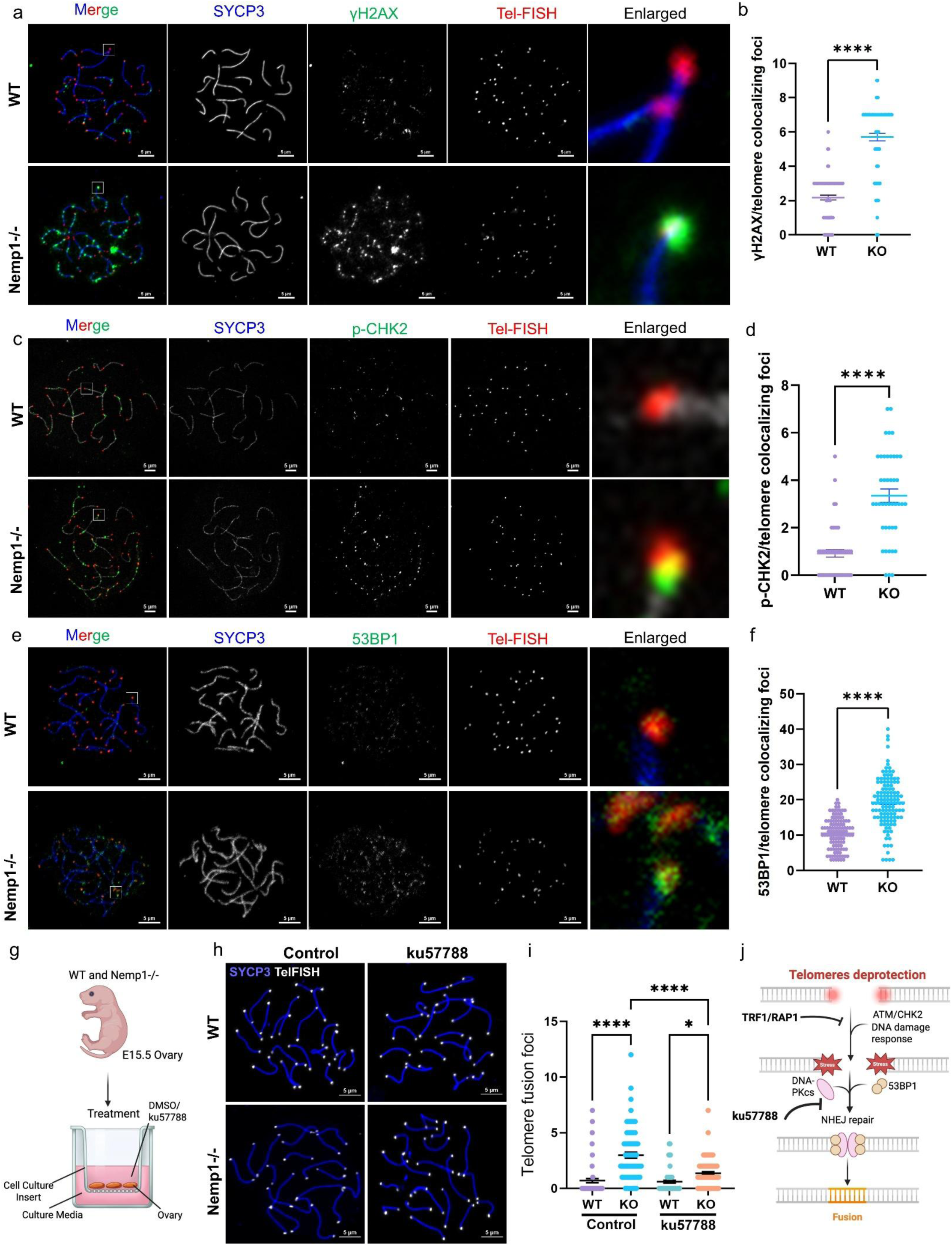
NEMP1 protects meiotic telomeres from persistent DNA damage and aberrant NHEJ-mediated fusion. (A) Representative images of pachytene-stage WT and *Nemp1*⁻/⁻ meiocytes showing telomeric DNA damage, detected by immunofluorescence staining for phosphorylated H2AX (γH2AX) and telomere FISH. DNA was counterstained with DAPI. Arrowheads indicate γH2AX-positive telomeres. (B) Quantification of the percentage of telomeres associated with γH2AX foci in WT and *Nemp1*⁻/⁻ pachytene meiocytes. (C) Representative images of phosphorylated CHK2 (pCHK2) and telomere FISH in WT and *Nemp1*⁻/⁻ pachytene meiocytes. Arrowheads indicate pCHK2-positive telomeres. (D) Quantification of telomere-associated pCHK2 signal intensity or the percentage of telomeres positive for pCHK2 in WT and *Nemp1*⁻/⁻ meiocytes. (E) Representative images showing 53BP1 localization at telomeres in WT and *Nemp1*⁻/⁻ pachytene meiocytes. Telomeres were detected by FISH, and 53BP1 was visualized by immunofluorescence. Arrowheads indicate telomere-associated 53BP1 foci. (F) Quantification of telomere-associated 53BP1 fluorescence intensity or the percentage of telomeres positive for 53BP1 in WT and *Nemp1*⁻/⁻ meiocytes. (G) Experimental scheme showing inhibition of classical nonhomologous end joining (c-NHEJ) using the DNA-PK inhibitor KU-57788 in WT and *Nemp1*⁻/⁻ meiocytes. (H) Representative images of telomere FISH in WT and *Nemp1*⁻/⁻ meiocytes treated with vehicle or KU-57788. Arrows indicate chromosome end-to-end fusions. (I) Quantification of telomere fusion frequency following KU57788 treatment. (J) Model illustrating the proposed mechanism by which NEMP1 protects meiotic telomeres from DNA damage and aberrant NHEJ-mediated chromosome-end fusion. Loss of NEMP1 leads to persistent telomeric DNA damage, increased 53BP1 recruitment, and aberrant NHEJ activity, resulting in telomere fusion and compromised chromosome-end integrity. Data are the mean ± s.d. of n = 3 independent trials per group (≥15 viable meiocyte spreads per trial). Statistical analyses were performed using two-sided unpaired t-tests (b, d, f). Statistical analyses were determined by one-way analysis of variance (ANOVA) followed by Tukey’s post hoc test (i). ****P < 0.0001. Scale bars, 5 μm. Schematic in g and j created in BioRender. (2026) https://BioRender.com/msdrrci.

Deprotected telomeres can be recognized as DNA breaks and undergo aberrant repair through classical non-homologous end joining (c-NHEJ), a process promoted by the DNA damage mediator 53BP1^61^. Deprotected telomeres are prone to aberrant repair through classical nonhomologous end joining (c-NHEJ), a process promoted by the DNA damage mediator 53BP1^61–63^. Consistent with this model, loss of NEMP1 in fetal and early postnatal ovaries resulted in robust recruitment of 53BP1 to telomeres, as evidenced by prominent 53BP1 colocalization at chromosome ends during meiotic prophase I in pachytene-stage meiocytes (Figure 5e, f).

To determine whether aberrant NHEJ contributes to the telomere fusions observed in *Nemp1* knockout meiocytes, we inhibited the NHEJ pathway using KU-57788 (Figure 5g). NHEJ inhibition partially suppressed telomere fusion in Nemp1 knockout meiocytes (Figure 5 h, i), indicating that aberrant NHEJ activity contributes to telomere instability and chromosome-end fusion in the absence of NEMP1 (Figure 5j). Together, these findings demonstrate that NEMP1 protects meiotic telomeres from persistent DNA damage and inappropriate NHEJ-mediated repair, thereby preserving chromosome-end integrity during meiotic prophase I.

### NEMP1 loss compromises telomere-led bouquet formation and rapid prophase movements

Our recent study showed that loss of *Nemp1* delays meiotic progression and causes persistent defects in chromosome synapsis that extend through pachytene ^37^. Consistent with these defects in meiotic chromosome organization, loss of NEMP1 markedly impaired telomere-led bouquet formation during prophase I (Figure 6a, b; Extended Data Figure 7a, b), and disrupted SYCP1 and SYCP3 organization (Extended Data Figure 7c, d). These findings suggested that NEMP1 may be required for the dynamic telomere movements that promote homolog pairing and chromosome organization during meiotic prophase I.

**Figure 6.**
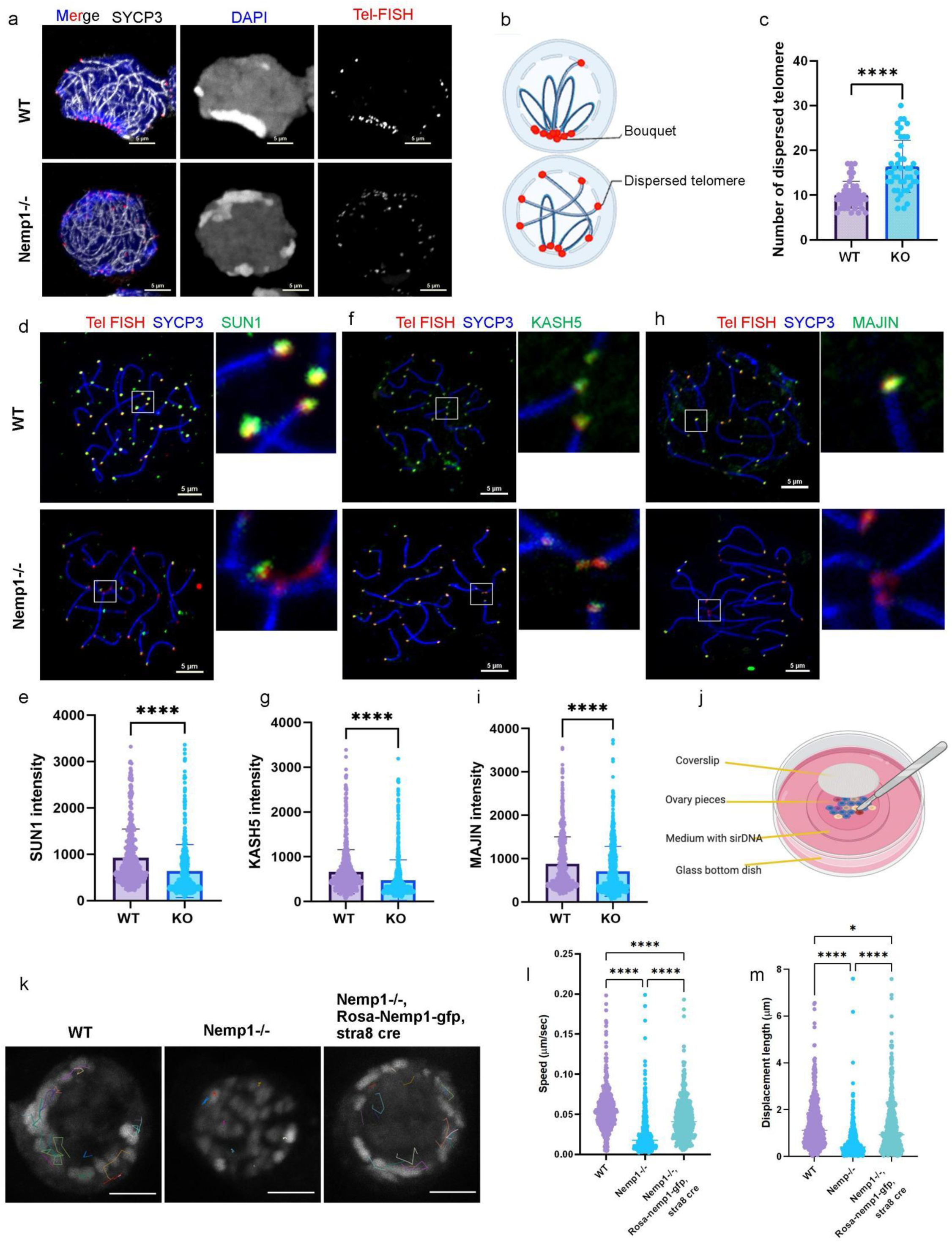
NEMP1 is required for telomere-led bouquet formation and rapid prophase movements. (A) Representative images of pachytene-stage WT and *Nemp1*⁻/⁻ meiocytes showing telomere organization and bouquet formation during meiotic prophase I. Telomeres were visualized by telomere FISH and chromosomes by SYCP3 immunostaining. (B) Quantification of telomere-led bouquet formation in WT and *Nemp1*⁻/⁻ meiocytes. (C) Representative images showing localization of SUN1 at meiotic telomeres in WT and *Nemp1*⁻/⁻ meiocytes. (D, E) Quantification of telomere-associated SUN1 signal in WT and *Nemp1*⁻/⁻ meiocytes. (F) Representative images of KASH5 and MAJIN localization at meiotic telomeres in WT and Nemp1⁻/⁻ meiocytes. (G–I) Quantification of telomere-associated KASH5 and MAJIN fluorescence intensity or localization in WT and *Nemp1*⁻/⁻ meiocytes. (J) Schematic of the live three-dimensional time-lapse imaging approach used to quantify rapid prophase movements (RPMs) in leptotene/zygotene meiocytes. Pericentromeric chromatin was labeled with SiR-DNA and tracked over time. (K) Representative time-lapse images and trajectories of chromatin movement in WT and *Nemp1*⁻/⁻ meiocytes. Arrowheads indicate tracked chromatin foci. (L) Quantification of RPM velocity in WT and *Nemp1*⁻/⁻ meiocytes. (M) Quantification of displacement length, defined as the cumulative distance traveled by tracked chromatin during the imaging period. (N) Representative live-cell imaging of *Nemp1*⁻/⁻ meiocytes expressing NEMP1-GFP following *Stra8*^P2Acre^ -mediated re-expression. (O, P) Quantification of RPM velocity and displacement following NEMP1-GFP re-expression compared with WT and *Nemp1*⁻/⁻ controls. Data are the mean ± s.d. of n = 3 independent trials per group (≥15 viable meiocytes per trial). Statistical analyses were performed using two-sided unpaired t-tests (c, e, g, i). Statistical analyses were determined by one-way analysis of variance (ANOVA) followed by Tukey’s post hoc test (l, m). *P < 0.05; ****P < 0.0001. Scale bars, 5 μm. Schematic in b and j created in BioRender. (2026) https://BioRender.com/msdrrci.

Meiotic telomere movement depends on SUN-domain proteins, which anchor chromosome ends to the nuclear envelope and couple meiotic chromosomes to cytoskeletal forces through LINC complexes ^12,16,64–69^. This coupling drives telomere movement and rotation along the nuclear envelope, thereby shuffling chromosomes and facilitating homolog recognition and pairing^70^. We therefore examined whether loss of NEMP1 affects the organization of the meiotic telomere– nuclear envelope machinery. Strikingly, Nemp1 knockout meiocytes showed significantly reduced association of SUN1, KASH5, and MAJIN with meiotic telomeres compared with wildtype controls (Figure 6 d-i). These findings suggested that NEMP1 deficiency compromises the organization or stability of the machinery required for telomere–nuclear envelope coupling.

To directly determine whether NEMP1 is required for meiotic chromosome movements, we adapted a live-cell three-dimensional time-lapse (4D) imaging approach in fetal ovary to monitor chromosome dynamics in prophase I meiocytes (Figure 6 j). Using SiR-DNA labeling, we tracked the movement of pericentromeric heterochromatin during meiotic prophase and correlated these dynamics with chromosome and synaptonemal complex organization, as previously described ^16^. Four-dimensional imaging revealed prominent rapid prophase movements (RPMs) in wild-type leptotene/zygotene meiocytes, characterized by coordinated chromatin movements and nuclear rotation (Figure 6k; Supplementary Video 7). In striking contrast, *Nemp1* knockout meiocytes exhibited significantly decrease of chromosome movement (Figure 6k; Supplementary Video 8). Quantitative tracking further demonstrated a profound reduction in both movement velocity and displacement length—the cumulative distance traveled by tracked chromatin over the imaging period—in *Nemp1* knockout meiocytes compared with wild-type controls (Figure 6l, m). Re-expression of NEMP1-GFP in *Nemp1*-deficient meiocytes using *Stra8*^P2Acre^ substantially restored rapid prophase chromosome movements, including movement velocity and displacement (Figure 6l, m; Supplementary Video 9). Thus, the loss of RPMs is a direct consequence of NEMP1 deficiency. To determine if the movements detected by our imaging approach reflect bona fide LINC-dependent chromosome dynamics, we analyzed meiocytes expressing dominant-negative KASH (DN-KASH)^71^ under *Stra8*^P2Acre^ control^71^. DNKASH expression caused a profound loss of RPMs, with meiocyte nuclei exhibiting virtually no residual rotation or coordinated chromosome movements (Extended Data Fig. 7e–g; Supplementary Video 10). The more severe phenotype observed following DN-KASH expression further supports the LINC dependence of the movements measured by our assay and places NEMP1 within the machinery that enables meiotic chromosome–nuclear envelope coupling.

Together, these findings establish NEMP1 as an essential regulator of meiotic chromosome dynamics. NEMP1 loss disrupts telomere-led bouquet formation and reduces the association of SUN1, KASH5 and MAJIN with meiotic telomeres, accompanied by a near-complete loss of LINC-dependent rapid prophase chromosome movements. Restoration of NEMP1 is sufficient to rescue these chromosome dynamics, demonstrating a direct functional requirement for NEMP1 in the machinery that couples meiotic chromosomes to nuclear envelope–cytoskeletal forces.

### NEMP1 associates with meiotic telomeres and the nuclear envelope–LINC machinery

The functional defects in chromosome movement and the reduced association of SUN1, KASH5, and MAJIN with meiotic telomeres in Nemp1 knockout meiocytes led us to investigate how NEMP1 is molecularly connected to the meiotic telomere–nuclear envelope machinery. We first examined the spatial relationship between NEMP1 and meiotic telomeres by immunofluorescence combined with telomere FISH. NEMP1 showed prominent enrichment at the meiocyte nuclear envelope and substantial colocalization with telomeric signals during meiotic prophase I (Figure 7 a), supporting a close association between NEMP1 and meiotic telomeres. To verify the association of NEMP1 with telomeric DNA, we performed a DNA pulldown assay (Figure 7 b). NEMP1 was enriched on TTAGGG repeats but not on the scrambled control sequence (Figure 7 c). Established telomere-associated proteins, including TRF1, TRF2 and RAP1, were also recovered in the telomere pulldown (Figure 7 c), validating the specificity of the assay. These findings indicate that NEMP1 associates with telomeric DNA, either directly or through a telomere-associated protein complex.

**Figure 7.**
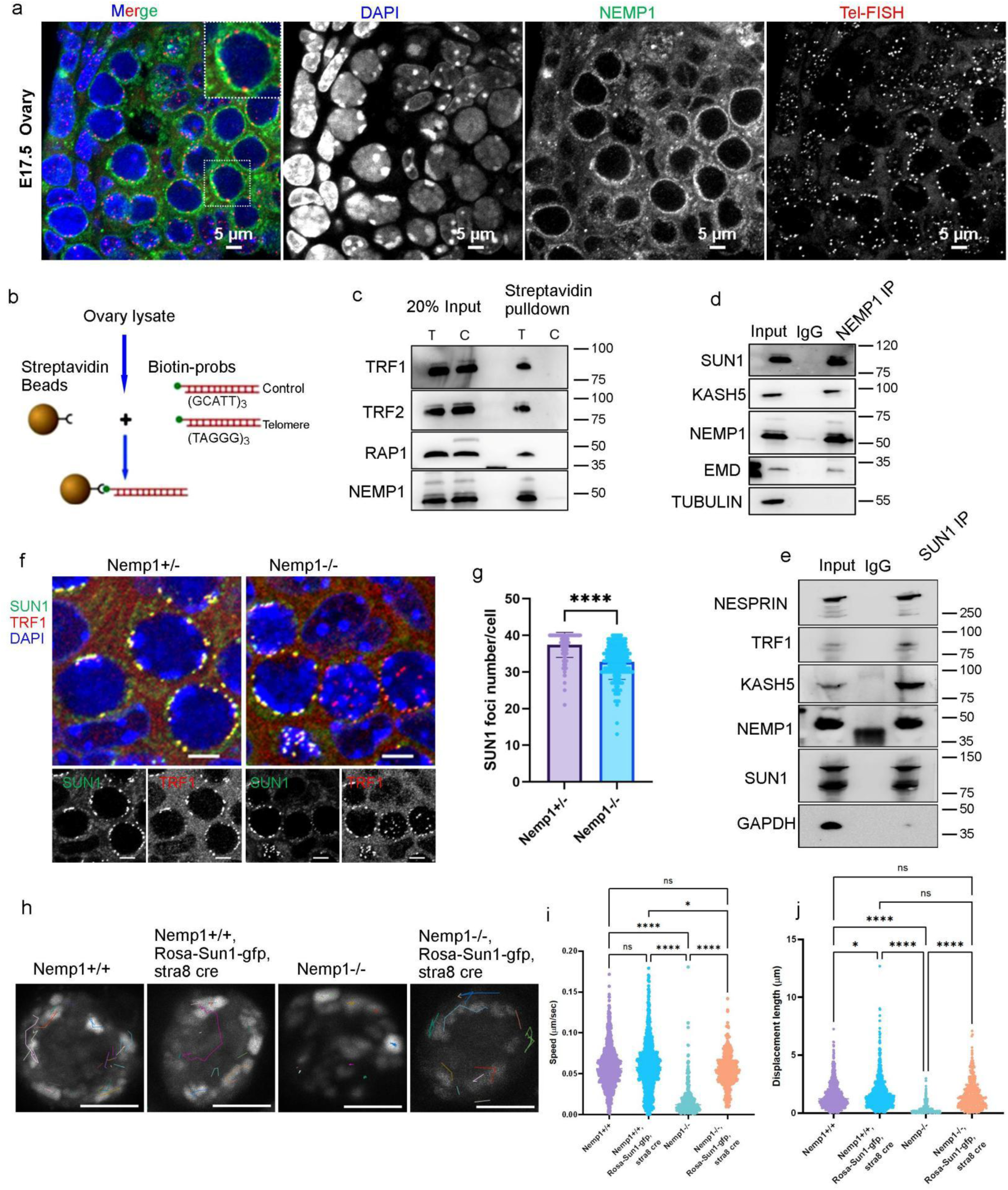
NEMP1 associates with meiotic telomeres and the nuclear envelope–LINC machinery. (a) Representative immunofluorescence images showing NEMP1 localization relative to telomeres in meiotic prophase I. NEMP1 was detected by immunofluorescence and telomeres by telomere FISH. Arrowheads indicate NEMP1-associated telomeres. (b) Schematic of the biotinylated telomere DNA pulldown assay used to examine the association of NEMP1 with telomeric chromatin. (c) Representative immunoblot analysis of NEMP1, TRF1, and RAP1 in telomeric DNA pulldown and control DNA pulldown fractions. Thirty ovaries from 15 E17.5 fetuses were pooled for the assay. (d) Co-immunoprecipitation of NEMP1 from E7.5 ovary extracts showing association with EMD, SUN1, and KASH5. IgG immunoprecipitation serves as a negative control. (e) Reciprocal coimmunoprecipitation showing recovery of NEMP1 in SUN1 immunoprecipitates. Thirty ovaries from 15 E17.5 fetuses were pooled for the assay. (f) Representative three-dimensional immunofluorescence images of E16.5 ovaries stained for SUN1 and the telomere-associated protein TRF1 in *Nemp1*⁺/⁻ and *Nemp1*⁻/⁻ meiocytes. Arrowheads indicate TRF1-positive telomeres. (g) Quantification of SUN1 fluorescence intensity at TRF1positive telomeres in *Nemp1*⁺/⁻ and *Nemp1*⁻/⁻ meiocytes. Data are the mean ± s.d. of n = 3 independent ovaries from 3 mice. (h) Representative live three-dimensional imaging of *Nemp1*⁻/⁻ meiocytes expressing SUN1-GFP following *Stra8*^P2Acre^-mediated re-expression. Representative trajectories of tracked chromatin foci are shown. (i, j) Quantification of rapid prophase movement velocity and displacement in WT, SUN1-GFP*-Stra8*^P2Acr^*, Nemp1*⁻/⁻, and Nemp1⁻/⁻; SUN1-GFP*-Stra8*^P2Acr^. Data are the mean ± s.d. of n = 3 independent trials per group (≥15 viable oocytes per trial). Statistical analyses were performed using two-sided unpaired t-tests (f). Statistical analyses were determined by one-way analysis of variance (ANOVA) followed by Tukey’s post hoc test (i, j). ****P < 0.0001. Schematic in b created in BioRender. (2026) https://BioRender.com/msdrrci.

**Figure 8.**
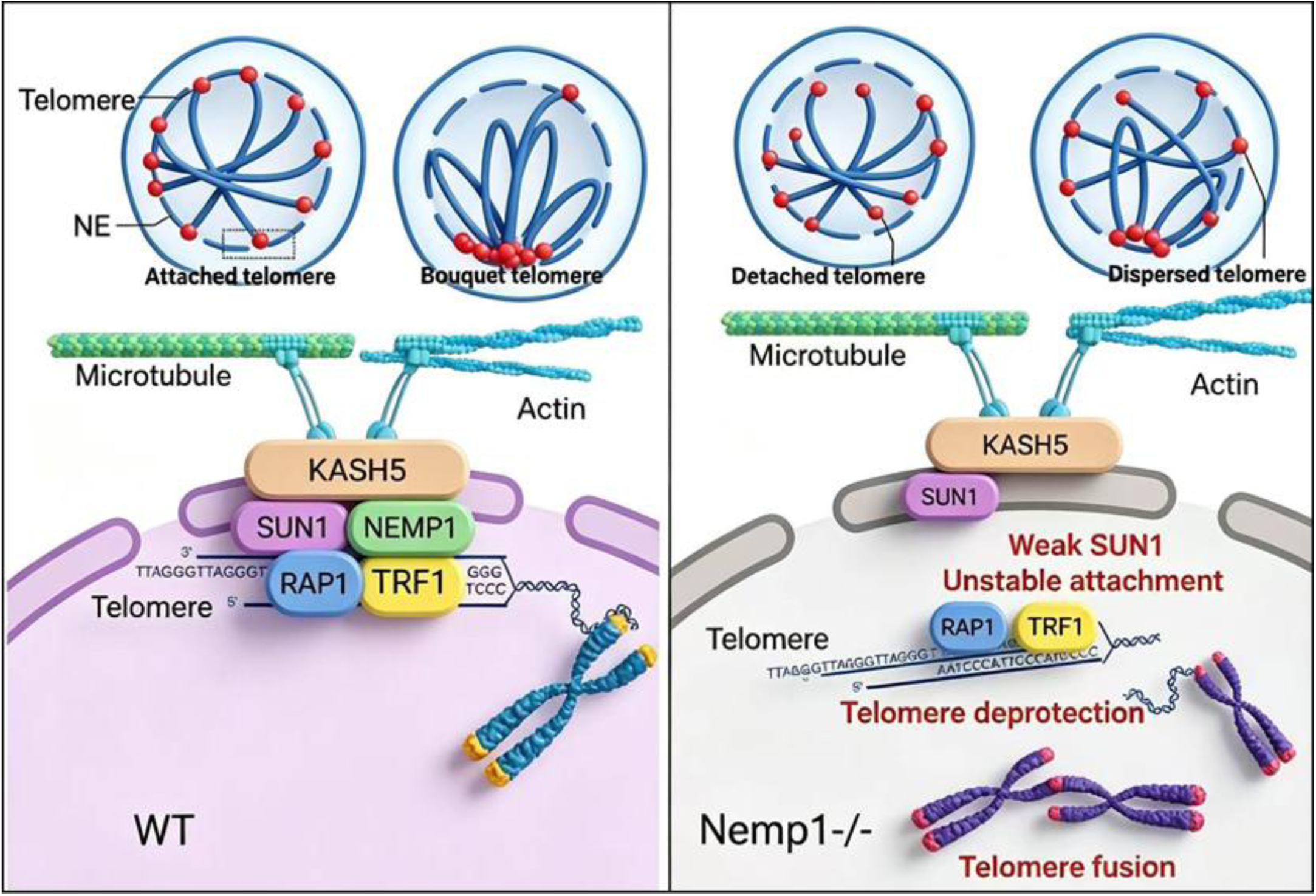
NEMP1 coordinates meiotic telomere–nuclear envelope coupling, chromosome dynamics, and telomere protection. During meiotic prophase I, NEMP1 localizes to the inner nuclear membrane and associates with telomeres and the SUN1-containing LINC machinery, promoting telomere attachment to the nuclear envelope, SUN1 recruitment, telomere-led bouquet formation, and rapid prophase chromosome movements (RPMs). This spatial and mechanical organization supports homolog pairing and synapsis while maintaining telomere integrity and protecting chromosome ends from persistent DNA damage and inappropriate repair. Loss of NEMP1 disrupts telomere–nuclear envelope coupling, reduces SUN1 recruitment, impairs bouquet formation and RPMs, and promotes telomere aggregation and progressive telomere shortening. Dysfunctional telomeres accumulate DNA damage and recruit 53BP1, leading to aberrant non-homologous end joining (NHEJ), chromosome end-to-end fusion, spindle and chromosome segregation defects, and ultimately aneuploidy and impaired female fertility. The graphical summary was created in BioRender. (2026) https://BioRender.com/msdrrci.

To define the molecular environment surrounding NEMP1 in the fetal ovary, we performed NEMP1-miniTurbo proximity labeling followed by quantitative proteomic analysis (Extended Data Fig. 8 a-d). The NEMP1 proximity proteome was significantly enriched for nuclear envelope- and LINC-associated proteins, including SUN1, KASH5, EMD and NESPRIN-2, together with SUMO1 (Extended Data Fig. 8c, d; Supplementary Table 1). Notably, NEMP1 was also identified in proximity to SUMO-associated machinery in an independent proximity-proteomics dataset^72^, providing orthogonal support for an association between NEMP1 and SUMO-enriched nuclear protein networks. In contrast, canonical components of the meiotic telomere complex, including MAJIN, TERB1 and TERB2, were not significantly enriched.

These findings place NEMP1 within a nuclear envelope–LINC-associated molecular environment and suggest that NEMP1 is unlikely to function as a core component of the canonical TERB1–TERB2–MAJIN telomere complex.

We next validated candidate NEMP1-associated proteins by co-immunoprecipitation. Immunoprecipitation of NEMP1 from E17.5 ovaries recovered EMD, SUN1 and KASH5, consistent with their enrichment in the NEMP1 proximity proteome, whereas these proteins were not substantially detected in control IgG precipitates (Fig. 7d). Reciprocal immunoprecipitation further confirmed the association between NEMP1 and SUN1 (Fig. 7e). Together, these biochemical data support a physical association between NEMP1 and the SUN1-containing nuclear envelope–LINC machinery in fetal ovary.

We next asked whether NEMP1 contributes to the recruitment of SUN1 to meiotic telomeres in vivo. Three-dimensional immunofluorescence analysis of E16.5 ovaries using SUN1 and the telomere-associated protein TRF1 revealed a marked reduction in SUN1 enrichment at TRF1positive telomeres in *Nemp1* knockout meiocytes compared with WT controls (Fig. 7f, g; Supplementary Video 11–12). Thus, loss of NEMP1 disrupts the association of SUN1 with meiotic telomeres, providing in vivo evidence that NEMP1 contributes to the organization of the telomere–nuclear envelope interface during meiotic prophase I. To determine whether increasing SUN1 availability could compensate for the loss of NEMP1, we expressed SUN1-GFP in WT and *Nemp1*-deficient meiocytes using Stra8^P2A-Cre^. SUN1-GFP expression did not significantly alter rapid prophase chromosome movements in WT meiocytes, indicating that increased SUN1 abundance alone does not enhance chromosome dynamics. In contrast, SUN1-GFP expression substantially restored chromosome movements in *Nemp1*-deficient meiocytes, as assessed by live three-dimensional imaging and quantitative measurements of movement velocity and displacement (Fig. 7h–j; Supplementary Video 13–16). Thus, the chromosome-movement defect caused by NEMP1 loss can be functionally rescued by increasing SUN1 availability. Together with the reduced recruitment of SUN1 to meiotic telomeres in Nemp1-deficient cells, these results support a model in which NEMP1 promotes SUN1-dependent coupling of meiotic telomeres to the nuclear envelope to facilitate chromosome movement.

Together, these findings position NEMP1 at the interface between meiotic telomeric chromatin and the nuclear envelope–LINC machinery. NEMP1 associates with telomeric chromatin and the SUN1-containing nuclear envelope machinery, whereas NEMP1 loss reduces SUN1 recruitment to meiotic telomeres and disrupts rapid prophase chromosome movements. Restoration of SUN1 is sufficient to rescue chromosome movement in *Nemp1*-deficient meiocytes, providing functional evidence for a NEMP1–SUN1 relationship in meiotic chromosome dynamics. We propose that NEMP1 functions as a novel nuclear envelope organizer that promotes the coupling of meiotic telomeres to the LINC machinery, thereby enabling chromosome movement and contributing to telomere integrity during meiotic prophase I.

## Discussion

Oocytes must preserve genome integrity over exceptionally long periods of time, often spanning decades in mammals. Failure of this protection underlies age-associated infertility, aneuploidy, and early embryonic loss. Despite its importance, the molecular architecture that safeguards the oocyte genome has remained poorly defined. NEMP1 is highly expressed and enriched at the oocyte nuclear envelope, where it directly associates with telomeres during meiotic prophase. Loss of NEMP1 partly disrupts telomere-NE attachment, leading to abnormal clustering, accelerated shortening, and the appearance of free chromosome ends lacking proper shelterin protection. These telomere defects precede overt chromosome fusion, aneuploidy, and sterility in *Nemp1* knockout females.

Previous studies established that meiotic telomere attachment relies on partially redundant pathways involving shelterin, TERB complexes, LINC complex and KASH components ^9,17,18,28,69,73^. However, the persistence of telomere attachment in most genetic models suggested the existence of additional INM regulators. Importantly, our findings extend beyond fully grown oocytes to early meiotic stages. Our recent studies in Drosophila and mouse models revealed that loss of NEMP1 triggers ATM–CHK2–dependent DNA damage signaling and causes extensive oocyte loss during fetal life, reflecting defective meiotic prophase progression and impaired homolog pairing and synapsis ^37^. Many nuclear proteins specifically bind DNA double-strand breaks to protect the lesion, signal damage to cell, activate repair pathways and delay cell-cycle enable to repair DNA damage, or trigger apoptosis when damage is irreparable ^74–79^. Here, we discovered that NEMP1 localizes to meiotic telomeres, where it restrains ATM–CHK2 signaling at chromosome ends and prevents pathological telomere fusion. The presence of 53BP1 at meiotic telomeres in the absence of NEMP1 suggests that NEMP1 normally functions to insulate chromosome ends from non-homologous end joining pathways. This provides a mechanistic explanation for how telomeres remain protected during the dynamic chromosome movements of meiosis, a period particularly vulnerable to genome instability ^9,11^. NEMP1 fills this gap by acting as an inner nuclear membrane–based scaffold that couples cytoskeletal forces to the nuclear envelope while simultaneously protecting chromosome ends from inappropriate NHEJ during meiotic chromosome dynamics. This rapid telomere movement facilitates proper selection of meiotic programmed DSB repair pathways that safeguard genome integrity ^25^.

Telomere tethering at the nuclear periphery is essential for efficient DNA double strand break repair in subtelomeric region^80,81^. Mechanistically, we propose that NEMP1 acts as a molecular bridge between telomeric chromatin and the inner nuclear membrane by promoting the formation or stabilization of a cooperative complex involving SUN1 and shelterin components. This architecture provides a direct physical linkage between chromosome ends and the nuclear envelope, stabilizing telomere position and preventing inappropriate DNA repair.

Our findings also provide a conceptual advance by directly linking nuclear envelope architecture to telomere protection. NEMP1 depletion compromises the telomere protection machinery as early as the fetal stage, with telomere defects subsequently becoming more pronounced during meiotic prophase and the prolonged prophase arrest of oocyte development. Consistent with this progressive deterioration, *Nemp1* knockout GV oocytes contained a population of telomeres lacking detectable TRF1 or RAP1 signals, indicating the accumulation of partially or fully deprotected chromosome ends. These findings suggest that NEMP1-dependent nuclear envelope organization is required for the establishment of telomere protection during early oogenesis, and that once this protection is lost, telomere dysfunction persists and may lead to progressive damage throughout subsequent oocyte development. The identification of SUMO1 in the NEMP1 proximity proteome is particularly intriguing given the established functions of SUMOylation in nuclear architecture, chromosome segregation and telomere-associated DNA repair^82^. Moreover, SUMO enrichment at damaged telomeres can facilitate the recruitment and cooperation of DNA repair factors, including RAD52, RAD51AP1 and BLM^83,84^. Together with the independent detection of NEMP1 within SUMO-associated proximity networks^72^, these observations raise the possibility that SUMO-dependent regulation contributes to the ability of NEMP1 to coordinate nuclear-envelope organization and telomere integrity. Determining whether NEMP1 itself is SUMOylated, or instead organizes SUMO-modified proteins at the meiotic nuclear envelope, will be an important direction for future investigation. While deprotected telomeres are known to activate ATM/ATR signaling and recruit 53BP1^21,85,86^, our data show that NEMP1 normally limits this response at meiotic telomeres. This is particularly important in oocytes, which lack robust telomere damage checkpoints during meiotic arrest and are therefore uniquely vulnerable to accumulating genome instability^14^. Loss of NEMP1 may consequently permit inappropriate engagement of DNA repair pathways that are normally suppressed or tightly regulated during meiosis, including non-homologous end joining (NHEJ) and alternative end joining ^87–89^. Consistent with this model, inhibition of NHEJ partially reduced telomere fusion in *Nemp1*-deficient meiocytes, supporting a role for aberrant NHEJ activity in the chromosome-end instability caused by NEMP1 loss. Together, these findings highlight the importance of nuclear-envelope-mediated regulation of DNA repair pathway accessibility in protecting meiotic telomeres from inappropriate repair.

More broadly, our study establishes the nuclear envelope as an active guardian of genome stability in oocytes and identifies NEMP1 as a critical structural protector of meiotic telomeres. Telomere dysfunction and progressive telomere attrition are recognized as forms of DNA damage that can activate the DNA damage response (DDR), ultimately contributing to cellular senescence^90–92^. NEMP1 is expressed in oocytes across metazoan species, suggesting that its role in oocyte nuclear organization may be evolutionarily conserved. Our findings show that, in mouse oocytes, NEMP1 is required during meiotic prophase I for proper telomere–nuclear envelope association, SUN1 recruitment, and telomere-led chromosome movement. Loss of NEMP1 disrupts these processes and is accompanied by persistent telomere deprotection, chromosome end-to-end fusions, aneuploidy and impaired oocyte developmental competence, with important implications for understanding the mechanisms driving female reproductive aging^93,94^. Consistent with this idea, common genetic variants near NEMP1 are associated with earlier age at menopause in large human cohorts ^95,96^. Together, our data identify NEMP1 as a regulator of the meiotic telomere–LINC interface and show that disruption of this interface during early oogenesis has lasting consequences for telomere integrity and chromosome stability. Whether variation in NEMP1 expression or function similarly affects telomere organization and oocyte quality in humans remains an important question for future investigation.

## Methods

### Mouse strains and animal husbandry

All mouse animal experiments were conducted in compliance with protocols approved by the Institutional Animal Care and Use Committee (IACUC) office under protocol #24-0170 at Washington University in Saint Louis. All mice were maintained in a specific pathogen-free condition in a controlled environment with a 12/12 h light/dark cycle. C57BL/6J and Rosa*Sun1*-gfp (B6;129-Gt(ROSA)^26Sortm5(CAG-Sun1/sfGFP)Nat/J^; Strain #:021039) mice were purchased from Jackson Laboratory. Nemp1 mutant mice (*Nemp1*^em#(TCP)McNeill^) were generated as previously described ^34^. Conditional knockout of *Nemp1* was achieved by introduction of an oocyte -specific Cre line (Zp3-cre). For stage-specific re-expression of NEMP1 or SUN1 on Nemp1 deficient background was achieved by introduction of meiosis-specific Cre line (*Stra8*^P2Acre^) ^97^ mice were crossed with mice carrying the corresponding conditional transgene Rosa-*Nemp1*-gfp or Rosa-*Sun1*-gfp. For validation of LINC-dependent chromosome movement, mice expressing dominant-negative KASH (DN-KASH) ^71^ under *Stra8*^P2Acre^ control were analyzed. Female fetal and postnatal ovaries were collected at the indicated developmental stages. Embryonic age was determined by the presence of a vaginal plug, with embryonic day 0.5 (E0.5) defined as noon on the day a plug was detected. Ovaries were dissected in prewarmed M2 medium (Sigma-Aldrich, Cat#M7167) and processed immediately for immunofluorescence, chromosome spreads, electron microscopy, biochemical assays, or live-cell imaging as indicated.

### Telomere and centromeric repeat (CentB) fluorescence in situ hybridization (FISH)

Telomeres or centromere were detected by FISH using a fluorescently labeled telomeric probe complementary to the mouse telomeric repeat sequence (TTAGGG)_3_ (PNABIO, cat#F1006) or centromere PNA FISH probe (ATTCGTTGGAAACGGGA) (PNABIO, cat#F3004). Fixed meiotic cells or chromosome spreads were prepared as described above and incubated with the telomere probe under hybridization conditions optimized for detection of mouse telomeric repeats. Briefly, samples are mixed directly with 500 nM PNA FISH probe in PNA hybridization buffer (PNABIO, cat# PFB01), incubated at 82°C for 10 min and left at room temperature in the dark for 24 h to complete the hybridization. After hybridization, the slides were sequentially washed 10 minutes each at 40°C: once in 1× SSC, once in 0.5× SSC, and once in 0.1× SSC. Following stringent washing, samples were counterstained with DAPI, mounted in anti-fade mounting medium (Agilent Technologies, cat# GM30411-2). Images were taken on a Nikon Eclipse Ti2 inverted confocal laser microscope (Nikon) with the NIS-Elements software using a 100× objective. Telomere foci number, fluorescence intensity, and telomere-associated structures were quantified using ImageJ. Telomere FISH signals were considered absent when fluorescence intensity fell below the predefined detection threshold established from wild-type controls. For colocalization experiments, telomere FISH was combined with immunofluorescence staining for NEMP1, TRF1, RAP1, H2AX, 53BP1, SUN1, MAJIN, KASH5 or other indicated proteins.

### Quantitative PCR analysis of relative telomere length in oocytes

Relative telomere length was measured using a qPCR-based assay adapted from previously described method ^47^. For each biological sample, five oocytes were pooled for genomic DNA extraction using KAPA Express Extract kit (Fisher scientific, cat# 50-196-5299). Telomeric DNA was quantified by qPCR and normalized to the multicopy nuclear reference gene Rn18s, which provides sufficient amplification from the limited amount of DNA obtained from small numbers of oocytes.

qPCR was performed using TB Green® Premix Ex Taq™ II (Takara, cat# RR82LR) on an CXF96 Touch™ Real-Time PCR Detection System. Each sample was analyzed in technical duplicate for both telomere and Rn18s amplification. The The Rn18s primers were: forward, 5′-AGAAACGGCTACCACATCCAA-3′; reverse, 5′CCTGTATTGTTATTTTTCGTCACTACCT-3′. Telomere primers were: forward, 5′CGGTTTGTTTGGGTTTGGGTTTGGGTTTGGGTTTGGGTT-3′; reverse, 5′GGCTTGCCTTACCCTTACCCTTACCCTTACCCTTACCCT-3′.

The cycling conditions were 95 °C for 2 min, followed by 40 cycles of 95 °C for 15 s, 60 °C for 30 s and 72 °C for 30 s, followed by standard melt-curve analysis. Telomere and Rn18s reactions from each sample were run on the same plate, and a common calibrator DNA sample was included on each plate to minimize inter-plate variation. Relative telomere length was calculated using the 2^−ΔΔCt^ method, with ΔCt calculated as telomere Ct minus Rn18s Ct and ΔΔCt calculated relative to the calibrator sample.

### Mouse oocyte collection

Germinal vesicle (GV)-stage oocytes were collected from female mice following intraperitoneal injection of 5 IU pregnant mare serum gonadotropin (PMSG). At 46 h after PMSG administration, mice were euthanized and ovaries were collected. Ovarian follicles were punctured using a 30-gauge needle under a stereomicroscope to release GV oocytes. Oocyte collection was performed in M2 medium supplemented with 1 μM milrinone (Sigma-Aldrich; cat# M4659) to maintain meiotic arrest at the GV stage. Granulosa cells and other ovarian somatic cells were removed from the follicular contents using a 40-μm cell strainer. GV oocytes were identified based on morphology and the presence of an intact germinal vesicle and collected by mouth pipetting under a stereomicroscope. For in vitro maturation, GV oocytes were thoroughly washed in milrinone-free M2 medium to allow meiotic resumption and subsequently cultured in M16 medium (Sigma-Aldrich; cat# M7292) at 37 °C in the presence of 5% CO_2_. Oocytes from the indicated experimental groups were collected at the corresponding developmental stages and processed for subsequent analyses.

### Immunofluorescence staining

Ovaries were collected at the indicated developmental stages and fixed in 4% paraformaldehyde for 24 h at room temperature (RT). Whole mount staining of ovaries and optical clearing were performed as previously described ^37^. For oocytes staining, oocytes were fixed in 4% PFA (Electron Microscopy Sciences, Cat#15700) in PBS solution at RT. Oocytes were removed from the PFA and then washed three times for 10 min each in PBS with 0.01% Tween-20 (PBST). Permeabilization was achieved through 0.5% Triton X-100 in PBS solution for 20 min and subsequently washed three times, 10 min each in PBST. Embryos were then blocked in blocking solution of 10% donkey serum diluted in PBS for 1 h at room temperature. Antibodies were diluted in blocking solution and incubated overnight at 4 °C. Oocytes were washed again three times for 10 min in PBST before being placed in secondary antibody solution with antibodies and 4,6-diamidino-2-phenylindole (DAPI) (VectorLabs Cat #H-1200-10) diluted in blocking buffer. Embryos were incubated for 2 h in secondary antibodies and washed six times in 10-min PBST washes. Oocytes were then transferred to a microbubble of PBS on a glass-bottomed MatTek dish (MatTek Corporation, P35G-1.5-20 C) for downstream imaging.

### Metaphase oocyte spreads

Metaphase spreads of oocytes during meiotic maturation were prepared from fully grown oocytes collected in M2 medium containing 1µM milrinone. Oocytes were matured in M16 medium at 37 °C in the presence of 5% CO_2_ for 8 hours (MI) or 16 h (MII). The zona pellucida was removed from oocytes using acidified Tyrode’s solution (Sigma-Aldrich, Cat# T1788). Oocytes were washed twice in M2 medium and transferred into drops of chromosome-spreading fixative containing 1% paraformaldehyde, 0.15% Triton X-100, and 3 mM dithiothreitol (DTT; pH 9.2). Oocytes were gently spread on glass slides and allowed to air-dry slowly at room temperature. After drying, chromosome spreads were subjected to telomere and centromeric repeat (CentB) fluorescence in situ hybridization (FISH) as described above.

### Meiotic chromosome spreads

Ovaries were collected at the indicated developmental stages and processed to generate meiotic chromosome spreads using standard hypotonic swelling and fixation procedures as previously described ^98^. Briefly, meiocytes were released into 100 mM sucrose in 5 mM sodium borate buffer (pH 8.5) by puncturing the ovaries with 29-gauge syringe needles and spread onto a glass-bottomed MatTek dish (MatTek Corporation, P35G-1.5-20 C) containing 2% paraformaldehyde, 0.3% Triton X-100, and 6 mM dithiothreitol (DTT; pH 9.2). Samples were allowed to air dry, rinsed with ultra-pure water and were subsequently processed for immunofluorescence and/or telomere FISH. Synaptonemal complex proteins were detected using antibodies against SYCP3 and, where indicated, SYCP1. Telomeres were visualized by telomere FISH. For quantitative analysis, identical acquisition settings were used for samples within each experiment. Telomere-associated fluorescence was quantified from individual telomeric foci or telomere-associated regions after background subtraction. For TRF1, RAP1, SUN1, KASH5 and MAJIN recruitment analysis, fluorescence intensity was measured at telomere FISH-positive telomeric foci in threedimensional image stacks. Quantification was performed using ImageJ software.

### Antibodies

The following antibodies were used; rabbit polyclonal antibody against NEMP1 (1:1000 for IF staining) ^34^, TMEM194 (Proteintech, cat#24477-1-AP) SUN1 (Abcam, cat# AB323868), γH2AX (Abcam,cat#ab11174), SCP1 (Abcam, cat#ab15090), Anti-CCDC155 (KASH5, Millipore, cat# HPA019940-100U), Anti-α-Tubulin−FITC (Sigma-Aldrich, cat#F2168); MAJIN (gift from Miguel A Brieño-Enríquez), SCP3 (Abcam, cat#ab314752), RAP1 (gift from de Lange Laboratory), TRF1 (gift from de Lange Laboratory), anti-DDX4 (Abcam, cat#ab13840), antipChk2 (p-T68) (Bioworld Technology, BS4043), anti-53BP1(Novus, cat# NB100-304) GAPDH (Sigma-Aldrich, cat#g9545); mouse polyclonal antibodies against SCP3 (Abcam, cat#ab97672), and γH2AX (Millipore, cat#05-636-I), Anti-α-Tubulin (Sigma-Aldrich, cat#T9026) and lamin A/C (Cell Signaling). Secondary antibodies used were Alexa Fluor goat anti-mouse 647 (1:100, Invitrogen), Alexa Fluor goat anti-mouse 405 (1:300, Invitrogen), Alexa Fluor goat anti-rabbit 488 (1:300, Invitrogen), Alexa Fluor donkey anti-rabbit 594 (1:300, Invitrogen). Streptavidin, Alexa Fluor™ 647 Conjugate (ThermoFisher, cat#S32357). Antibodies were diluted 1:500 for immunostaining and 1:1000 for western blotting.

### Three-dimensional imaging and analysis of telomere–nuclear envelope association

For three-dimensional analysis of meiotic telomeres, optical z-stacks were acquired through the entire nucleus using a Nikon Eclipse Ti2 inverted confocal laser microscope (Nikon) with the NIS-Elements software using a 100× objective. Z-step size and acquisition parameters were kept constant between experimental groups. Three-dimensional reconstruction and quantitative analysis were performed using Imaris. Telomeres were identified by telomere FISH. Telomeres were classified as nuclear-envelope associated or internalized based on their spatial relationship with the nuclear envelope. The proportion of internalized telomeres was quantified from individual meiocytes across multiple animals.

### Analysis of meiotic bouquet formation

Bouquet formation was assessed in leptotene, zygotene, and pachytene meiocytes based on the spatial clustering of telomeres at one region of the nuclear envelope. Telomeres were detected by telomere FISH, chromosomes/synaptonemal complexes by SYCP3 immunostaining and DAPI. For each meiocyte, the spatial distribution of telomeres was assessed using three-dimensional image stacks. Cells were classified as bouquet-positive when telomeres exhibited a concentrated distribution over a defined region of the nuclear envelope according to predefined criteria. Quantification was performed blind to genotype.

### Live-cell imaging of meiotic chromosome movements

A three-dimensional time-lapse imaging method was developed to quantify rapid prophase movements (RPMs) in living meiocytes. Ovaries were isolated at the indicated developmental stage and maintained in M16 medium supplemented with 10 nM SiR-DNA (cytoskeleton, cat#CY-SC007) under controlled temperature, humidity, and CO conditions. To avoid the rotation of the meiocytes in the medium, dishes were pre-treated with Cell-Tak (Sigma-Aldrich, cat#CLS354240) for 30 min before cell spreading. Meiocytes were placed on glass-bottom dishes and gently covered with a coverslip to minimize cell movement and prevent floating during live-cell imaging. Meiocytes were identified based on nuclear morphology and meiotic stage. Three-dimensional image stacks were acquired every 30 s for 10 min using a ×100 objective on a Nikon microscope equipped with NIS-Elements software. Exposure times were 0.1 s for GFP and 0.07 s for SiR-DNA. Pericentromeric hetrochromatin foci were tracked through successive three-dimensional frames using Image J. RPM velocity was calculated from the displacement of individual tracked chromatin foci over time. Displacement length was defined as the cumulative distance traveled by each tracked chromatin focus during the imaging period. All genotypes were imaged using identical acquisition parameters whenever possible.

To validate that the chromosome movements measured by live imaging depend on functional LINC-mediated nuclear envelope–cytoskeletal coupling, meiocytes expressing dominant-negative KASH under *Stra8*^P2Acre^ control were analyzed. RPMs were measured using the same three-dimensional time-lapse imaging and tracking procedure described above. Chromosome movement velocity and displacement length were compared between control and DN-KASH-expressing meiocytes. For NEMP1 rescue experiments, NEMP1-GFP was re-expressed in Nemp1-deficient meiocytes using the *Stra8*^P2Acre^ system. Meiocytes from Nemp1-deficient animals carrying the NEMP1-GFP rescue allele were identified by genotype and GFP fluorescence. For SUN1 rescue experiments, SUN1-GFP was re-expressed in Nemp1-deficient meiocytes using *Stra8*^P2Acre^. SUN1-GFP-positive meiocytes were identified by GFP fluorescence and analyzed using live three-dimensional time-lapse microscopy. RPM velocity, displacement length, and nuclear rotation were quantified using the same tracking pipeline used for NEMP1 rescue experiments. All groups were analyzed using matched imaging and tracking parameters.

### Electron microscopy analysis of meiotic telomeres

Ovaries E17.5 were processed for transmission electron microscopy using the indicated fixation, dehydration, embedding, and sectioning procedures. Testes were isolated, dissected and incubated in a fixative consisting of 2.5% glutaraldehyde plus 4% paraformaldehyde in PBS, for 2 hr at RT. Samples were washed three times for 30 min at RT in sodium cacodylate buffer then post-fixated in 2% OsO4 in sodium cacodylate buffer for 1 hr at 4°C. Samples were washed in sodium cacodylate buffer at 4°C for 3 x 15 min. Pre-embedding staining was done overnight at RT in the dark using 0.5% uranyl acetate. Then samples were washed with utra-purify water for 2 x 15 min. Samples were then dehydrated in an ethanol series, 2 x 5 min for each concentration of ethanol: 50%, 70%, 80%, 90%, 95%, 100%. For plastic embedding, samples were treated with 100% propylene oxide for 2 x 10 min at RT then incubated in 1:1 propylene oxide:Durcupan (epoxic resin durcupan ACM) for at least 1 hour with cap, then left overnight without cap and in desiccators. Samples were then embedded in Durcupan and left for 3-6 hr without cap in desiccators at RT.

Resin polymerization was done at 60°C for at least 48 hr. Sections of 95 nm were cut with a Leica Ultracut UCT ultramicrotome (Leica Microsystems Inc., Bannockburn, IL), stained with uranyl acetate and lead citrate, and viewed on a JEOL 1200 EX transmission electron microscope (JEOL USA Inc., Peabody, MA) equipped with an AMT 8 megapixel digital camera and AMT Image Capture Engine V602 software (Advanced Microscopy Techniques, Woburn, MA). Meiotic telomere ultrastructure was assessed based on the presence of electron-dense structures associated with the ends of synapsed lateral elements and their attachment to the inner nuclear membrane.

### Detection of telomeric DNA damage

Telomeric DNA damage was assessed by combined immunofluorescence for DNA damage markers and telomere FISH. Meiotic spreads were stained for γH2AX, phosphorylated CHK2, or other indicated DNA damage markers followed by telomere FISH. Telomere-associated DNA damage was quantified based on the colocalization of DNA damage foci with telomeric FISH signals. The number and fluorescence intensity of telomere-associated damage foci were quantified using Image J.

### 53BP1 recruitment to telomeres

Meiotic chromosome spreads were immunostained for 53BP1 followed by telomere FISH. Three-dimensional image stacks were acquired under identical conditions between genotypes.

53BP1 fluorescence associated with individual telomeres was quantified after background subtraction. The fraction of telomeres positive for 53BP1 and the fluorescence intensity of telomere-associated 53BP1 were determined for individual meiocytes.

### NHEJ inhibition

To examine the contribution of classical non-homologous end joining to telomere fusion, ovaries were treated with the DNA-PK inhibitor KU-57788 (MCE, cat#HY-11006) 1 μM for 24h and change fresh medium for 24 h. Control samples received the corresponding vehicle. Following treatment, meiotic chromosome spreads were prepared and analyzed by telomere FISH and immunofluorescence. Chromosome end-to-end fusion frequency and telomere-associated abnormalities were quantified as described above.

### Telomere DNA pulldown

Telomeric DNA pulldown assays were performed to determine whether NEMP1 associates with telomeric DNA, as previously described^63^. Briefly, biotinylated telomeric DNA containing tandem mouse telomeric repeats (TTAGGG)n (PNABIO, cat#F2002) was prepared and immobilized on streptavidin-conjugated magnetic beads (ThermoFischer, cat#88816). A corresponding non-telomeric DNA (GCATT)n (PNABIO) was used as a control. Protein extracts from 30 ovaries were incubated with the immobilized telomeric DNA for 4h at 4°C with rotation. Beads were extensively washed with PBS to remove nonspecific proteins. Bound proteins were eluted in SDS sample buffer and analyzed by immunoblotting. NEMP1 was detected using an anti-NEMP1 antibody. TRF1, TRF2 and RAP1 were analyzed as positive controls for telomere-associated proteins.

### NEMP1-TurboID proximity labeling

To identify proteins in proximity to NEMP1 in vivo, we performed miniTurbo-based proximity labeling using mouse fetal ovarian tissue. A Rosa26-driven *Nemp1*–miniTurbo mouse line was crossed with *Stra8*^P2Acre^ mice to induce expression of the NEMP1–miniTurbo fusion protein in the germ cell lineage. Timed matings were established, with the morning of vaginal plug detection designated embryonic day 0.5 (E0.5).

Pregnant females carrying the appropriate embryos were administered biotin beginning at E10.5 to enable in vivo proximity-dependent biotinylation. Biotin (Sigma-Aldrich, cat#B4501) was administered for 7 days, and fetal ovaries were collected at E17.5. Control ovaries lacking the NEMP1–miniTurbo fusion protein were processed in parallel to distinguish specific NEMP1proximal proteins from endogenously biotinylated or nonspecifically enriched proteins. Following dissection, fetal ovaries were collected in ice-cold phosphate-buffered saline (PBS) and lysed in NP40 lysis buffer (Invitrogen, cat# J60766.AP) containing 1% protease inhibitor cocktail (Invitrogen, cat#87786). Tissue lysates were homogenized and clarified by centrifugation. Biotinylated proteins were enriched using streptavidin-conjugated beads. After extensive washing to remove nonspecifically bound proteins, enriched proteins were subjected to on-bead digestion with trypsin.

Resulting peptides were analyzed by liquid chromatography–tandem mass spectrometry (LC– MS/MS) by the Washington University in St. Louis Mass Spectrometry (MassSpec) Team. Protein identification and quantitative analysis were performed by comparison with the appropriate mouse protein database. Proteins enriched in NEMP1–miniTurbo samples relative to control samples were considered candidate NEMP1-proximal proteins. Differential enrichment was evaluated using quantitative proteomic analysis, and significantly enriched proteins were further examined for functional associations with the nuclear envelope, LINC complex, telomere organization, and meiotic chromosome dynamics. This approach identified several candidate proteins associated with NEMP1, including components of the nuclear envelope and meiotic chromosome movement machinery, such as SUN1, KASH5, and Nesprin-2, supporting the presence of an NEMP1-associated protein network at the nuclear envelope.

### Co-immunoprecipitation

For co-immunoprecipitation experiments, E17.5 ovaries were collected and lysed in NP40 lysis buffer (Invitrogen, cat# J60766.AP) containing 1% protease inhibitor cocktail (Invitrogen, cat#87786). Lysates were clarified by centrifugation. Equal amounts of protein (20 ovaries) were incubated with an antibody against NEMP1, SUN1, or the indicated protein overnight at 4°C with rotation. Protein G/A magnetic beads (Invitrogen, cat# 88802) were added and incubated for an additional 2 h. Parallel samples incubated with control IgG were processed in parallel.

Beads were washed extensively with lysis buffer and bound proteins were eluted in SDS sample buffer (BIO-RAD, cat#1610737). Immunoprecipitated proteins were resolved by SDS–PAGE and analyzed by immunoblotting using antibodies against NEMP1, SUN1, KASH5, EMD, or other indicated proteins. For reciprocal co-immunoprecipitation, SUN1 immunoprecipitation was performed and the presence of NEMP1 was assessed by immunoblotting.

### Immunoblotting

Protein samples were separated by SDS–PAGE and transferred onto PVDF or nitrocellulose membranes. Membranes were blocked with 5% nonfat milk in TBST and incubated with primary antibodies overnight at 4°C. After washing, membranes were incubated with HRP-conjugated secondary antibodies and visualized using a Bio-Rad chemiluminescence imaging system. Protein abundance was quantified and normalized to the indicated loading control or input.

### Statistical analysis

All experiments were performed using independent biological replicates as indicated in the figure legends. Individual meiocytes were treated as the experimental unit unless otherwise specified. Data are presented as mean ± SEM unless otherwise indicated.

Statistical analyses were performed using GraphPad Prism 11. For comparisons between two groups, an unpaired two-tailed Student’s t-test was used. For comparisons involving multiple groups, one-way ANOVA followed by the indicated multiple-comparison test was used. A P value < 0.05 was considered statistically significant. Statistical details, including the exact test, n values, and definition of biological replicates, are provided in the corresponding figure legends and source data.

## Acknowledgements

We thank the Mouse Genetics Core at Washington University School of Medicine in St. Louis, particularly Dr. Mia Wallace, for mouse husbandry and genotyping. We thank Molecular Microbiology Imaging Facility and Dr. Wandy L Beatty for ovary transmission electron microscopy. The Mass Spectrometry Technology Access Center at the McDonnell Genome Institute MTAC@MGI-Washington University School of for proteomics. We are grateful to Miguel Angel Brieño-Enríquez, Luis Batista, Chun-Kan Chen, Huanyu Qiao, and Titia de Lange for helpful comments and discussions. We also thank Dr. Titia de Lange, Dr. Miguel Angel Brieño-Enríquez for providing antibodies. The graphical summary was created with BioRender.com. We thank the members of the McNeill laboratory for helpful discussions and feedback.

## Funding

This work is supported by the National Institute of Child Health and Human Development (NICHD), United States (Grant RO1-HD108639 to H.M.); The Lalor Foundation fellowship (to H. Z.) and Postdoctoral Fellow Seed of Independence Grant (to H. Z.).

## Supplementary Video Legends

**Supplementary Video 1 | Chromosome segregation during meiotic maturation in wild-type oocytes.**

Representative time-lapse imaging of chromosome segregation in a wild-type oocyte during meiotic maturation. Chromosomes were visualized using SiR-DNA. The movie shows chromosome separation during the metaphase-to-anaphase transition. Related to Fig. 1.

**Supplementary Video 2 | Defective chromosome segregation during meiotic maturation in *Nemp1*−/− oocytes.**

Representative time-lapse imaging of an *Nemp1*−/− oocyte during meiotic maturation, with chromosomes visualized using SiR-DNA. The *Nemp1*−/− oocyte exhibits defective chromosome segregation during the metaphase-to-anaphase transition compared with wild-type oocytes. Related to Fig. 1.

**Supplementary Video 3 | Three-dimensional organization of telomeres in a wild-type GV oocyte.**

Representative three-dimensional reconstruction of telomere FISH in a wild-type germinal vesicle (GV)-stage oocyte. Telomeres are distributed throughout the nucleus as discrete foci, with no prominent telomere aggregates. Related to Fig. 2 a, b.

**Supplementary Video 4 | Telomere aggregation in a Nemp1−/− GV oocyte.**

Representative three-dimensional reconstruction of telomere FISH in an *Nemp1*−/− GV-stage oocyte. Multiple telomere signals form prominent aggregates, demonstrating abnormal telomere organization following NEMP1 loss. Related to Fig. 2 a, b.

**Supplementary Video 5 | Three-dimensional organization of telomeres and TRF1 in a wildtype GV oocyte.**

Representative three-dimensional reconstruction of telomere FISH and TRF1 immunofluorescence in a wild-type GV-stage oocyte. Telomeres are present as discrete nuclear foci and are associated with the telomere-protection protein TRF1. Related to Fig. 2 g, f.

**Supplementary Video 6 | Telomere aggregation and loss of TRF1 association in a *Nemp1*−/− GV oocyte.**

Representative three-dimensional reconstruction of telomere FISH and TRF1 immunofluorescence in an *Nemp1*−/− GV-stage oocyte. Telomere aggregates are evident, and a subset of telomeric signals shows reduced or undetectable TRF1 signal, consistent with impaired telomere protection following NEMP1 loss. Related to Fig. 2 g, f.

**Supplementary Video 7 | Telomere-led chromosome movements in wild-type meiocytes.** Representative three-dimensional time-lapse imaging of a wild-type meiocyte during meiotic prophase I. Telomere-associated chromosome ends undergo dynamic movements along the nuclear periphery, illustrating rapid telomere-led chromosome movements. Related to Fig. 6 k-m.

**Supplementary Video 8 | Impaired telomere-led chromosome movements in *Nemp1*−/− meiocytes.**

Representative three-dimensional time-lapse imaging of an *Nemp1*−/− meiocyte during meiotic prophase I. Telomere-led chromosome movements are markedly reduced compared with wildtype meiocytes, demonstrating impaired chromosome dynamics following NEMP1 loss. Related to Fig. 6 k-m.

**Supplementary Video 9 | NEMP1-GFP restores telomere-led chromosome movements in Nemp1−/− meiocytes.**

Representative three-dimensional time-lapse imaging of an *Nemp1*−/− meiocyte expressing NEMP1-GFP using *Stra8*-P2A-Cre. NEMP1-GFP expression restores telomere-led chromosome movements along the nuclear periphery, supporting a requirement for NEMP1 in normal meiotic chromosome dynamics. Related to Fig. 6 k-m.

**Supplementary Video 10 | DN-KASH expression disrupts rapid prophase chromosome movements.**

Representative three-dimensional time-lapse imaging of meiocytes expressing dominantnegative KASH (DN-KASH) during meiotic prophase I. DN-KASH expression causes a profound loss of rapid prophase chromosome movements (RPMs), consistent with disruption of LINC complex-dependent coupling between meiotic telomeres and cytoskeletal forces. Related to Extend Fig. 6 e-g.

**Supplementary Video 11 | Three-dimensional organization of SUN1 at meiotic telomeres in WT meiocytes.**

Representative three-dimensional reconstruction of SUN1 and TRF1 immunofluorescence in an E16.5 WT meiocyte. SUN1 is enriched at TRF1-positive telomeres at the nuclear periphery, consistent with the formation of meiotic telomere–nuclear envelope attachment sites. Related to Fig. 7f, g.

**Supplementary Video 12 | Reduced SUN1 enrichment at meiotic telomeres in Nemp1−/− meiocytes.**

Representative three-dimensional reconstruction of SUN1 and TRF1 immunofluorescence in an E16.5 *Nemp1*−/− meiocyte. Compared with WT controls, SUN1 enrichment at TRF1-positive telomeres is markedly reduced, demonstrating impaired organization of the telomere–SUN1 interface following NEMP1 loss. Related to Fig. 7f,g.

**Supplementary Video 13 | Rapid prophase chromosome movements in wild-type meiocytes.** Representative three-dimensional time-lapse imaging of chromosome movements in wild-type meiocytes during meiotic prophase I. Chromosome ends exhibit robust rapid prophase movements along the nuclear periphery. Related to Fig. 7h–j.

**Supplementary Video 14 | Rapid prophase chromosome movements in wild-type meiocytes expressing SUN1-GFP.**

Representative three-dimensional time-lapse imaging of wild-type meiocytes expressing SUN1GFP. Robust chromosome movements are maintained following SUN1-GFP expression, indicating that increased SUN1 availability does not substantially perturb normal chromosome dynamics. Related to Fig. 7h–j.

**Supplementary Video 15 Impaired rapid prophase chromosome movements in *Nemp1*−/− meiocytes.**

Representative three-dimensional time-lapse imaging of chromosome movements in *Nemp1*−/− meiocytes during meiotic prophase I. Chromosome movements show reduced velocity and displacement compared with wild-type controls, demonstrating defective rapid prophase chromosome movements following NEMP1 loss. Related to Fig. 7h–j.

**Supplementary Video 16 Restored chromosome dynamics following SUN1-GFP expression in *Nemp1*−/− meiocytes.**

Representative three-dimensional visualization of chromosome trajectories in *Nemp1*−/− meiocytes expressing SUN1-GFP. The reconstructed trajectories illustrate restoration of the spatial range and dynamics of chromosome movements following SUN1-GFP expression. Related to Fig. 7h–j.

---

**Figure S1.**
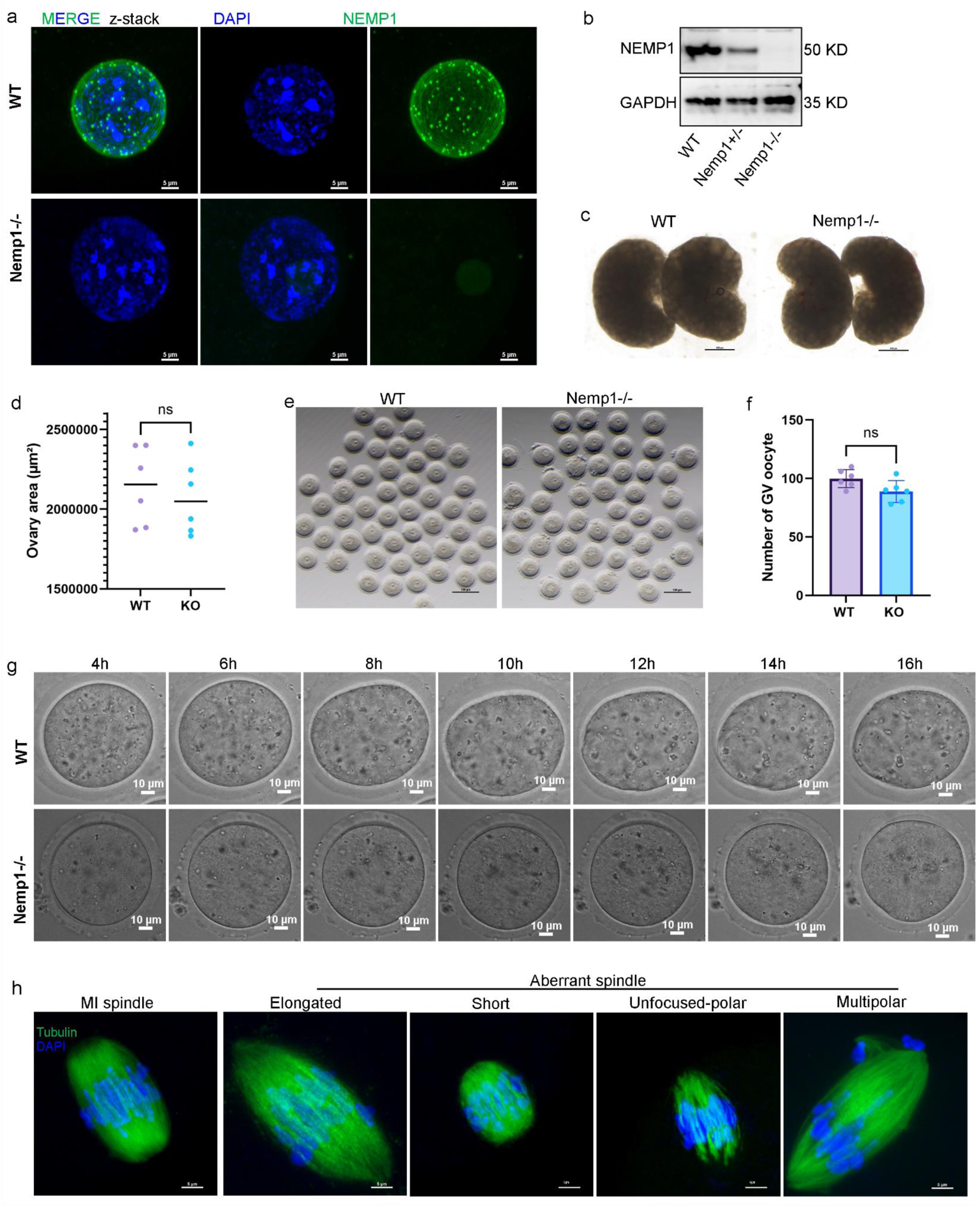
Validation of NEMP1 depletion and characterization of oocyte recovery and meiotic progression. (a) Representative immunofluorescence images showing NEMP1 localization in wild-type (WT) and Nemp1⁻/⁻ germinal vesicle (GV) oocytes from 21day mouse. In WT oocytes, NEMP1 is enriched at the nuclear envelope and forms prominent punctate foci, whereas NEMP1 signal is absent in *Nemp1*⁻/⁻ oocytes. Scale bar, 5 μm. (b) Western blot analysis of NEMP1 protein expression in whole ovaries from WT, *Nemp1*⁺/⁻, and *Nemp1*⁻/⁻ females. (c) Representative ovary images from WT and *Nemp1*⁻/⁻ mouse. n=6 ovaries from 3 mice. (d) Quantification of ovarian areas, showing loss of NEMP1 does not significantly affect ovary size. (e) Representative GV oocyte images from WT and *Nemp1*⁻/⁻ mouse. n=12 ovaries from 6 mice. (f) Quantification of oocyte recovery from WT and *Nemp1*⁻/⁻ females, showing that loss of NEMP1 does not significantly affect the number of oocytes recovered. (g) Time-lapse images of in vitro maturation of WT and *Nemp1*⁻/⁻ oocytes from 4 to 16 h. Representative images show that, whereas WT oocytes progressed through meiosis and extruded the first polar body, *Nemp1*⁻/⁻ oocytes frequently remained arrested at metaphase I (MI). Time after the initiation of in vitro maturation is indicated. Scale bar, 5μm. (h) Representative images of spindle morphology in WT and *Nemp1*⁻/⁻ metaphase I oocytes. *Nemp1*⁻/⁻ oocytes display increased frequencies of abnormal spindle morphologies, including elongated, shortened, multipolar, and unfocused spindles. Chromosomes and spindle microtubules are labeled as indicated. Scale bar, 5 μm..

**Extended Data Figure 2.**
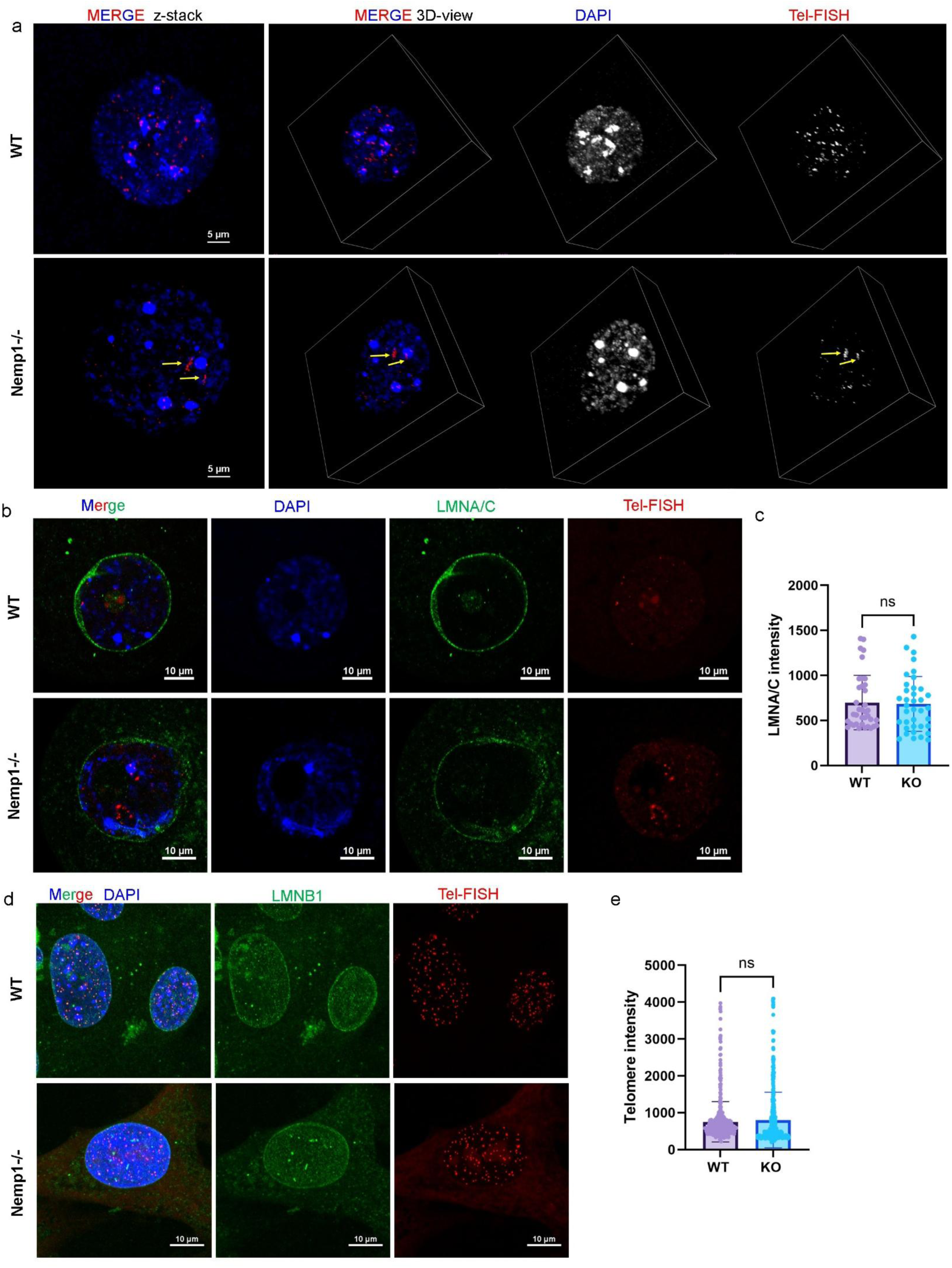
NEMP1 depletion disrupts telomere organization independently of overt nuclear lamina defects. (a), Representative three-dimensional (3D) Z-stack images of telomere FISH in control and *Nemp1*⁻/⁻ germinal vesicle (GV) oocytes. Control oocytes display discrete, spatially separated telomere foci throughout the nucleus, whereas *Nemp1*⁻/⁻ oocytes frequently exhibit prominent telomere aggregates (TAs), characterized by the coalescence of multiple telomeres into large foci. Enlarged views of representative nuclei are shown. Scale bars, 5 μm. (b), Representative immunofluorescence images showing lamin A/C distribution in control and *Nemp1*^⁻/⁻^ GV oocytes. Lamin A/C displays a comparable nuclear envelope-associated distribution in both genotypes. Scale bars, 10 μm. (c), Quantification of lamin A/C fluorescence intensity in control and *Nemp1*^⁻/⁻^ GV oocytes, showing no significant difference between genotypes. (d), Representative telomere FISH images of control and *Nemp1*^⁻/⁻^ mouse embryonic fibroblasts (MEFs). No obvious telomere aggregation phenotype is observed following NEMP1 depletion. Scale bars, 10 μm. (e), Quantification of fluorescence intensity in control and *Nemp1*^⁻/⁻^ MEFs, showing no significant differences between genotypes. Data are the mean ± s.d. of n = 3 independent. Each data point represents one oocyte/cell. Statistical analyses were performed using two-sided unpaired t-tests. Exact sample sizes, numbers of independent experiments, and P values provided in the corresponding source data.

**Extended Data Figure 3.**
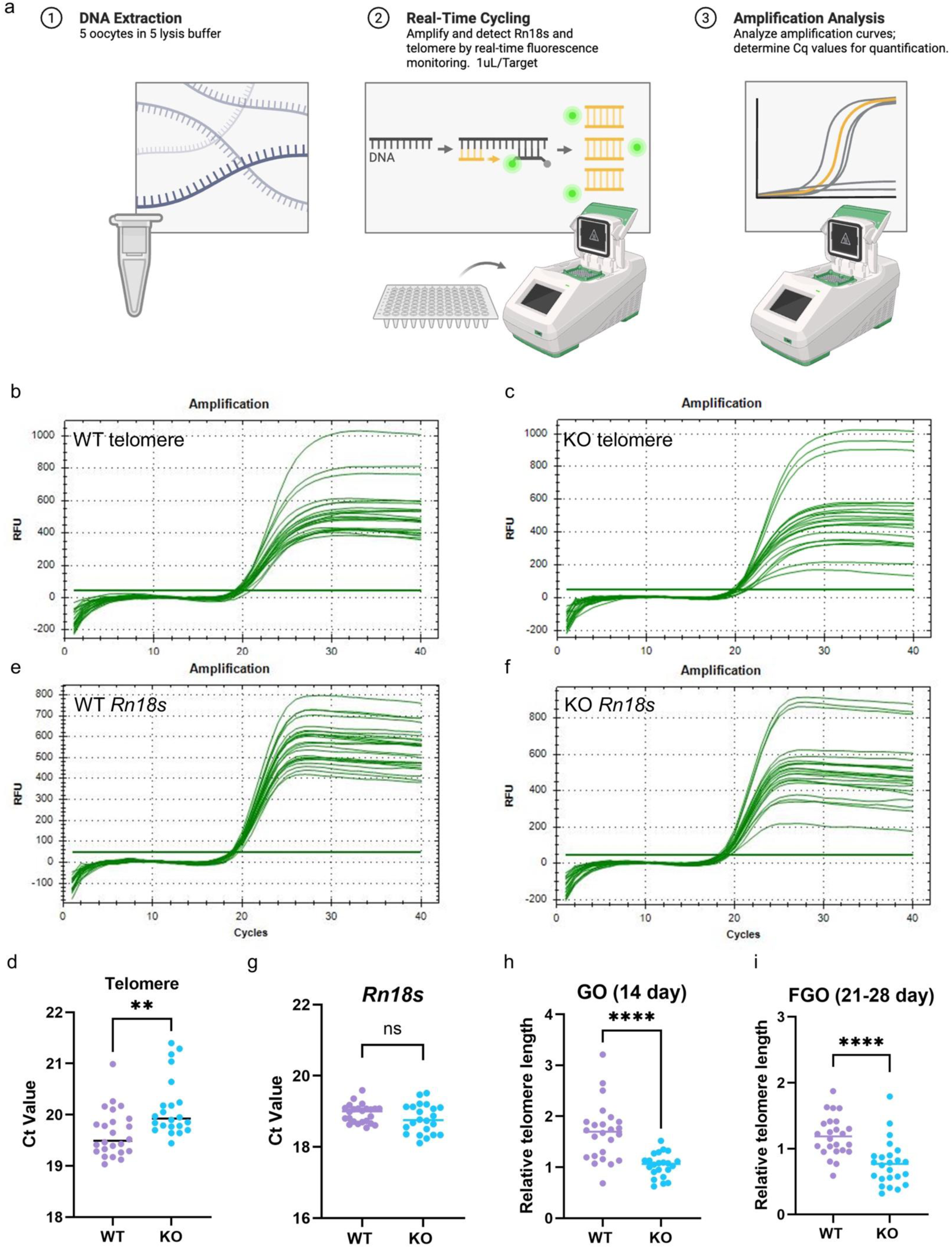
Quantification of relative telomere length in oocytes by qPCR. (a) Schematic of the qPCR-based assay used to quantify relative telomere length in oocytes. Telomeric DNA was quantified relative to the multicopy nuclear reference gene Rn18s. Five oocytes were pooled for each biological sample. Telomere DNA abundance was quantified relative to the nuclear reference gene Rn18s. (b, c, e, f) Representative qPCR amplification curves for telomere DNA and *Rn18s*, respectively, in individual growing oocytes (GO) and fully grown oocytes (FGO). (d, g) Representative cycle threshold (Ct) values for telomere DNA and Rn18s in oocytes, demonstrating reproducible amplification across samples and genotypes. (h, i) Relative telomere length in control and *Nemp1*^⁻/⁻^ GO (14 day) and FGO (21-28 day) oocytes, calculated from telomere DNA abundance normalized to *Rn18s*. *Nemp1*^⁻/⁻^ oocytes exhibit significantly reduced relative telomere length compared with control oocytes at both developmental stages. Data are the mean ± s.d. of n = 3 independent trials per. Each data point represents 5 oocytes. Statistical analyses were performed using two-sided unpaired t-tests. Exact sample sizes, numbers of independent experiments, and P values provided in the corresponding source data. Schematic in a created in BioRender. (2026) https://BioRender.com/msdrrci.

**Extended Data Figure 4.**
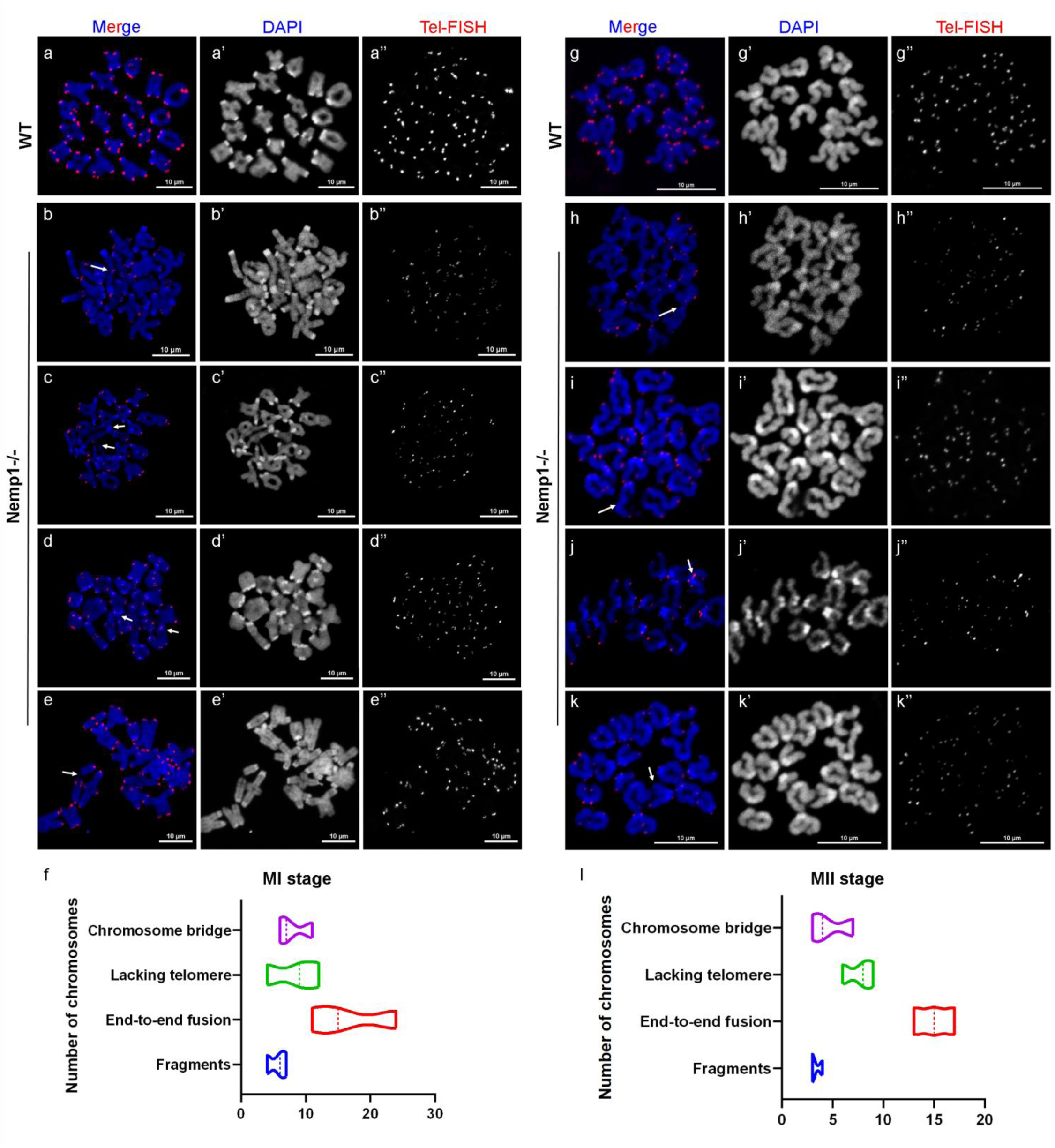
Chromosome abnormalities in NEMP1-deficient oocytes. (a-d) Representative metaphase I (MI) chromosome spreads from control and *Nemp1*^⁻/⁻^ oocytes showing chromosome morphology and telomeric DNA detected by FISH. (b–d), Representative images highlighting the major chromosome abnormalities observed in *Nemp1*⁻/⁻ oocytes, including chromosome bridges (b), chromosome ends lacking detectable telomeric signals (c), end-to-end chromosome fusions (d) and chromosome fragments (e). Scale bars, 10 μm. (f) The number of different types of chromosome defects in *Nemp1*^⁻/⁻^ oocytes. (h-k) Representative metaphase II (MII) chromosome spreads from control and *Nemp1*^⁻/⁻^ oocytes showing chromosome morphology and telomeric DNA detected by FISH. Representative images highlighting the major chromosome abnormalities observed in *Nemp1*⁻/⁻ oocytes, including chromosome bridges (h), chromosome fragments (i) end-to-end chromosome fusions (j) and chromosome ends lacking detectable telomeric signals (k). Scale bars, 10 μm. (l) The number of different types of MII chromosome defects in *Nemp1*⁻/⁻ oocytes.

**Extended Data Figure 5.**
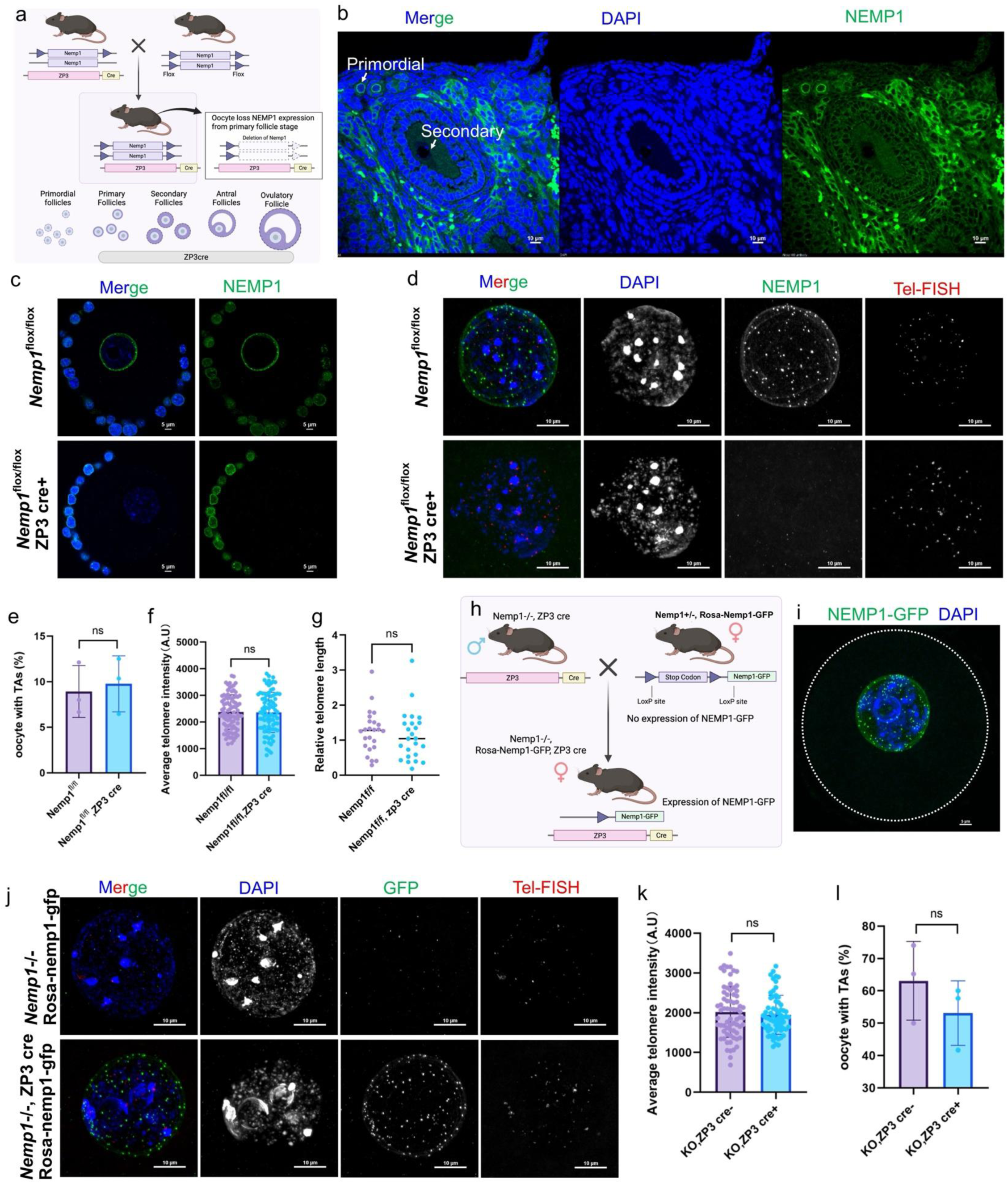
NEMP1 associates with telomeres and functions during early oogenesis. (a), Schematic of conditional *Nemp1* deletion using Zp3-Cre. (b) Immunofluorescence validation of NEMP1 loss at secondary follicular oocytes from *Nemp1*^flox/flox^; Zp3-Cre. (c) Representative immunofluorescence images validating conditional NEMP1 deletion in oocytes from Nemp1^flox/flox^; Zp3-Cre females. NEMP1 signal was absent in oocytes but retained in the surrounding cumulus cells, confirming oocyte-specific deletion of NEMP1 in this conditional knockout model. Scale bar, 5 μm. (d) Representative telomere FISH and NEMP1 immunofluorescence images. (e) Quantification of telomere aggregates in control and Zp3-Cre–deleted oocytes. (f, g) Telomere fluorescence intensity and relative length analysis in GV-stage oocytes from *Nemp1*^flox/flox^ (Control) and *Nemp1*^flox/flox^; Zp3-Cre females showing no detectable telomere shortening. (h) Schematic of reexpression of NEMP1 from a Rosa-*Nemp1*-GFP allele driven by Zp3-Cre. (i) Representative GFP fluorescence images validating NEMP1-GFP re-expression. (j) Representative telomere FISH and NEMP1-GFP fluorescence images from *Nemp1*⁻/⁻ and Rosa-Nemp1-GFP; Zp3-Cre in *Nemp1*⁻/⁻ background. (k) Quantification of telomere aggregates in *Nemp1*⁻/⁻ and NEMP1-GFP re-expressing. (l) Telomere fluorescence intensity analysis in GV-stage oocytes from *Nemp1*⁻/⁻ and NEMP1-GFP re-expressing GV oocyte. Re-expression of NEMP1 at the Zp3-Cre stage did not restore telomere organization or telomere fluorescence intensity in *Nemp1*⁻/⁻ oocytes. Data are the mean ± s.d. of n = 3 independent trials per group (≥15 viable oocytes per trial). Statistical analyses were performed using two-sided unpaired t-tests. Exact sample sizes, numbers of independent experiments, and P values provided in the corresponding source data. Schematic in h created in BioRender. (2026) https://BioRender.com/msdrrci.

**Extended Data Figure 6.**
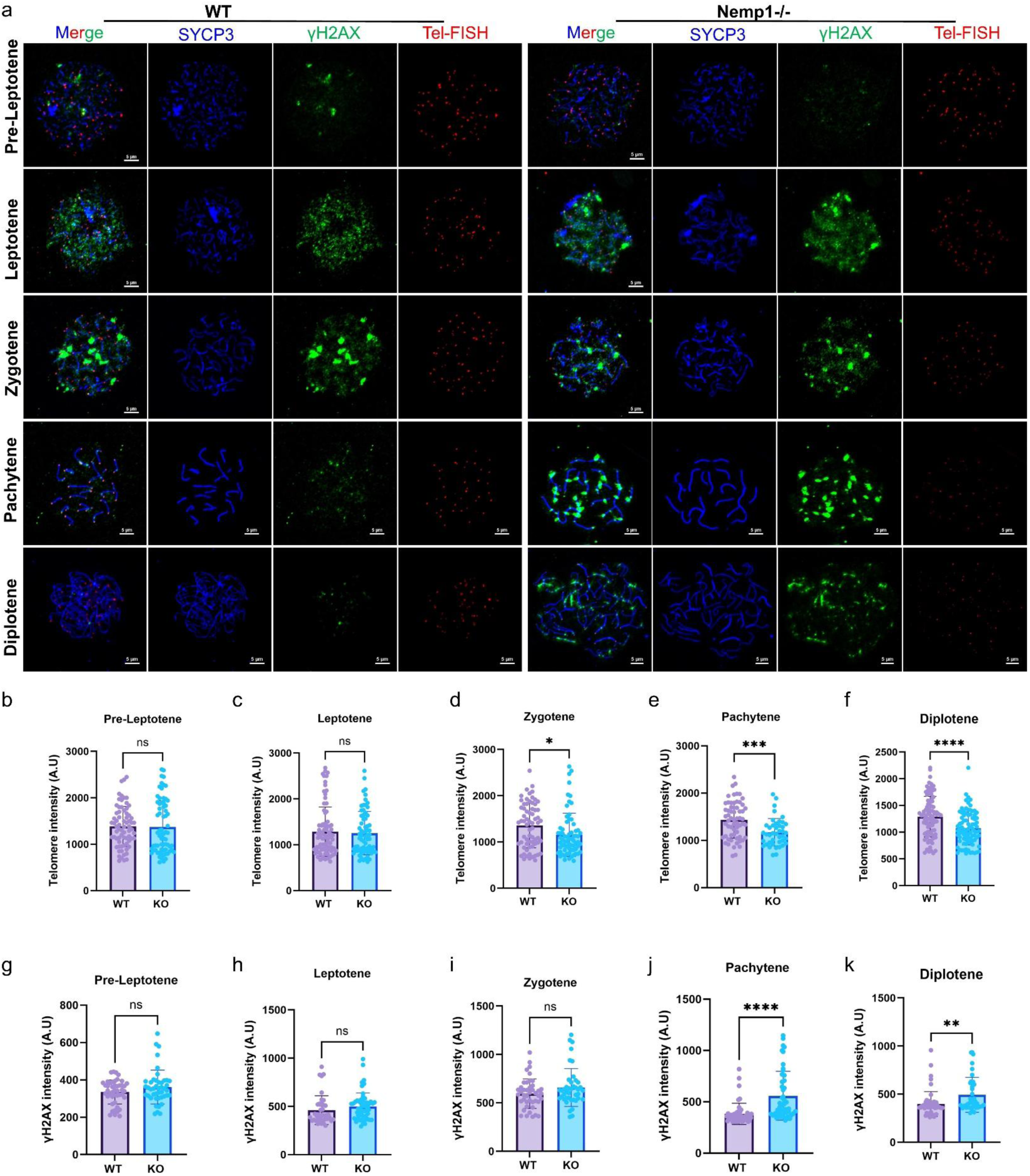
Telomere and DNA damage dynamics during meiotic prophase. **I.** (a). Chromosome spreads from E17.5 oocytes of WT and *Nemp1*⁻/⁻ females. Meiotic prophase I stages were identified based on SYCP3 morphology and γH2AX staining and included pre-leptotene, leptotene, zygotene, pachytene and diplotene stages. Scale bar, 5 μm. (b–f), Quantification of telomere fluorescence intensity in chromosome spreads from pre-leptotene (b), leptotene (c), zygotene (d), pachytene (e) and diplotene (f) oocytes from WT and *Nemp1*⁻/⁻ females. (g-h), Quantification of γH2AX fluorescence intensity in chromosome spreads from the indicated meiotic prophase I stages in WT and *Nemp1*⁻/⁻ oocytes. Data are the mean ± s.d. of n = 3 independent trials per group. Each data point represents an individual chromosome spread. Statistical analyses were performed using two-sided unpaired t-tests. Exact sample sizes, numbers of independent experiments, and P values provided in the corresponding source data.

**Extended Data Figure 7.**
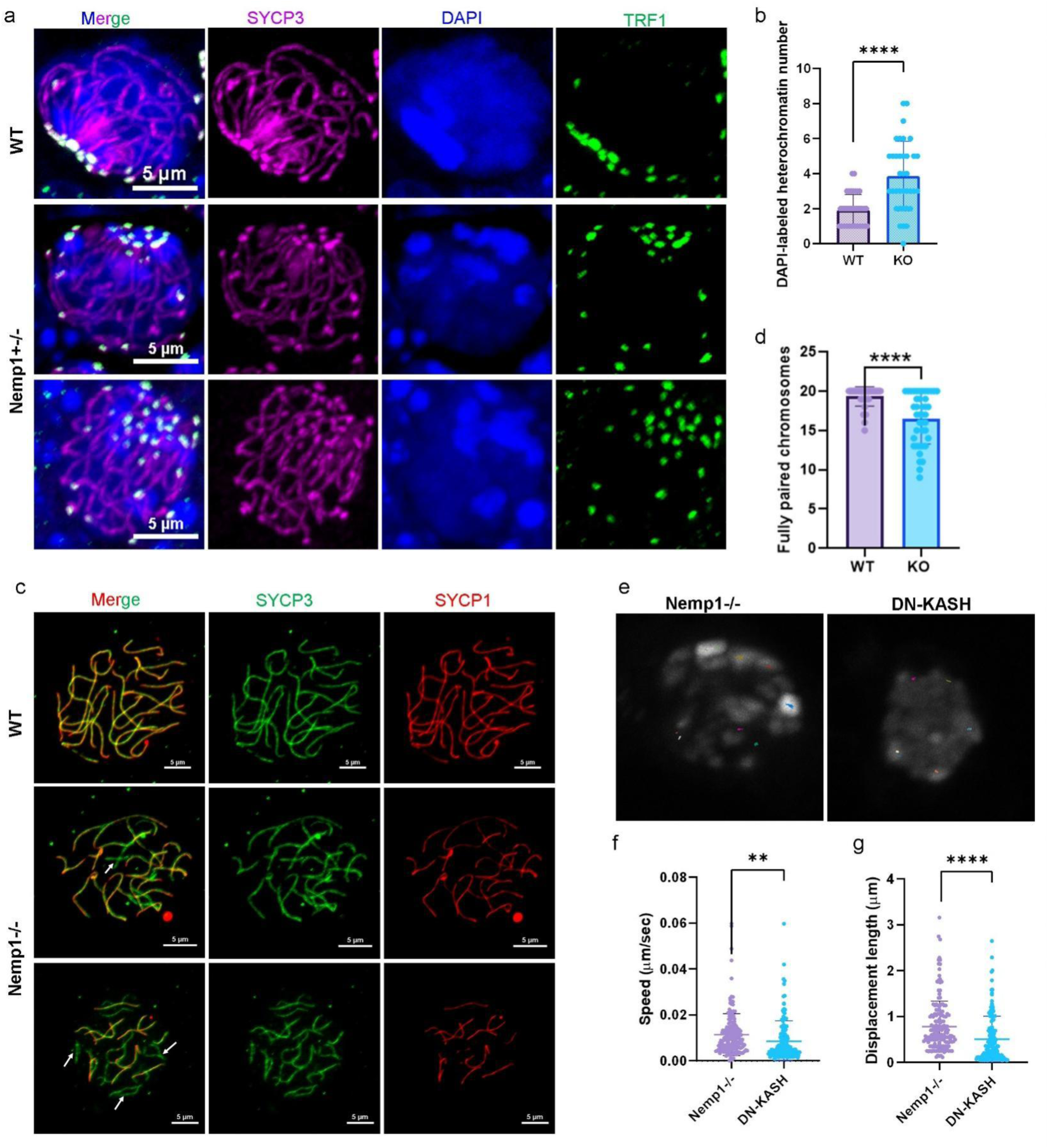
NEMP1 deficiency disrupts meiotic chromosome synapsis and LINC-dependent chromosome movements. (a,b) Representative images (a) and quantification (b) of telomere-led bouquet formation in E17.5 ovaries from WT and *Nemp1*⁻/⁻ meiocytes. *Nemp1*⁻/⁻ meiocytes show a reduced frequency of heterochromatin clustering at one side of the nuclear envelope, consistent with impaired bouquet formation. Scale bar, 5 μm. (c,d), Representative immunofluorescence images (c) and quantification (d) of SYCP1 and SYCP3 organization in meiotic prophase I chromosome spreads from WT and *Nemp1*⁻/⁻ meiocytes. Loss of NEMP1 results in disrupted synaptonemal complex organization and reduced chromosome synapsis. Scale bar, 5 μm. (e), Representative time-lapse images of meiotic prophase I nuclei expressing dominant-negative KASH (DN-KASH) under *Stra8*^P2Acre^ control. DN-KASH expression markedly reduces chromosome movements and nuclear rotation compared with control meiocytes. Time points are indicated. Scale bar, 5 μm. (f, g), Quantification of RPM velocity and displacement in *Nemp1*^-/^and DN-KASH-expressing meiocytes. DN-KASH expression results in a profound reduction in chromosome movement and nuclear rotation, with no detectable residual coordinated nuclear or chromatin movements. Data are the mean ± s.d. of n = 3 independent trials per group. Each data point represents an individual chromosome spread. Statistical analyses were performed using two-sided unpaired t-tests. **P < 0.01; ****P < 0.0001.

**Extended Data Figure 8.**
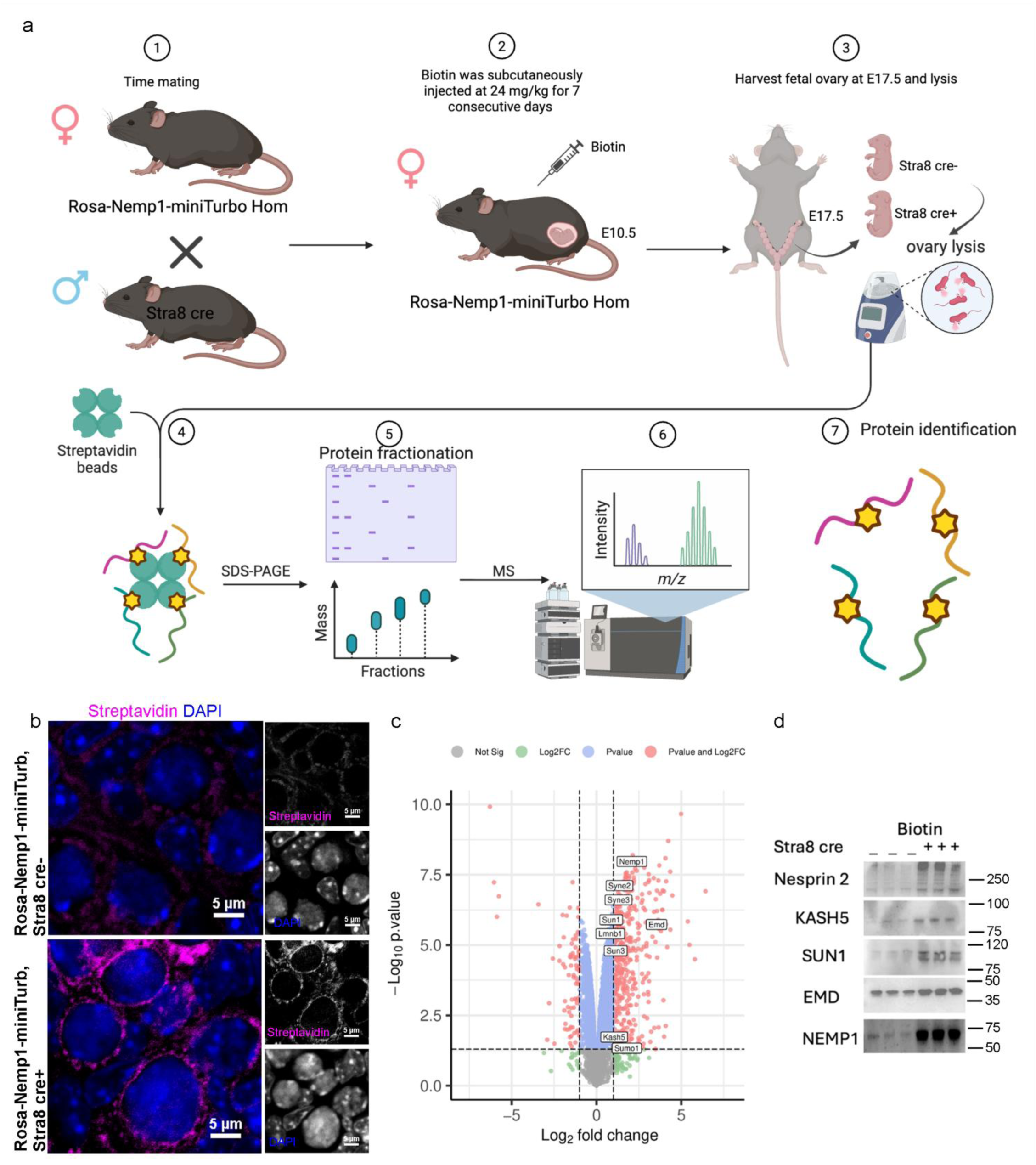
NEMP1-miniTurbo identifies a nuclear envelope and LINC-associated protein environment. (a), Schematic of the NEMP1-miniTurbo proximity-labeling strategy and experimental workflow. NEMP1TurboID was used to biotinylate proteins in the local molecular environment of NEMP1 at the nuclear envelope. (b) Validation of NEMP1-miniTurbo expression, localization and proximity labeling. Representative images showing NEMP1-miniTurbo localization and streptavidin staining to detect TurboID-dependent protein biotinylation in E17.5 ovary. (c) Volcano plot of the NEMP1-miniTurbo proximity proteome. Proteins enriched in the NEMP1-TurboID sample relative to the control are highlighted, including nuclear envelopeand LINCassociated proteins such as EMD, SUN1, KASH5 and Nesprin-2. (d), Immunoblot validation of selected proteins identified in the NEMP1-TurboID proximity proteome. Streptavidin pull-down of biotinylated proteins followed by immunoblotting demonstrates enrichment of EMD, SUN1, KASH5 and Nesprin-2 in the NEMP1miniTurbo sample compared with the control. Three independent biological replicates were performed. For each replicate, 12 ovaries from six E17.5 fetuses were pooled for the assay. Schematic in a created in BioRender. (2026) https://BioRender.com/msdrrci.

## Notes

### Competing Interest Statement

The authors have declared no competing interest.

